# A Bayesian network approach to assess the influence of climate change and pesticide use practices on the ecological risks of pesticides in a protected Mediterranean wetland

**DOI:** 10.1101/2022.12.15.520567

**Authors:** Claudia Martínez-Megías, Sophie Mentzel, Yasser Fuentes-Edfuf, S. Jannicke Moe, Andreu Rico

## Abstract

Pollution by agricultural pesticides is one of the most important pressures affecting Mediterranean coastal wetlands. Pesticide risks are expected to be influenced by climate change, which will result in an increase of temperatures and a decrease in annual precipitation rates in this region. On the other hand, pesticide dosages are expected to change given the increase in pest resistance and the implementation of environmental policies like the European ‘Farm-to-Fork’ strategy, which aims for a 50% reduction in pesticide usage by 2030. The influence of climate change and pesticide use practices on the ecological risks of pesticides needs to be evaluated making use of realistic environmental scenarios. This study aimed to assess how different climate change and pesticide use practices affect the ecological risks of pesticides in the Albufera Natural Park (Valencia, Spain), a protected Mediterranean coastal wetland. We performed a probabilistic risk assessment for nine pesticides applied in rice production using scenarios comprised of three climatic regimes (the 2008 record, and projections for 2050 and 2100), three pesticide application regimes (the recommended dose, and 50% increase and 50% decrease), and their combinations. The scenarios were used to simulate pesticide exposure concentrations in the water column of the rice paddies using the RICEWQ model. Pesticide effects were characterized using acute and chronic Species Sensitivity Distributions built with laboratory toxicity data for aquatic organisms. Risk quotients were calculated as probability distributions making use of a Bayesian network approach, and best fitting distributions for the calculated exposure data and the SSDs. Our results show that future climate projections will influence exposure concentrations for some of the studied pesticides, yielding higher dissipation and lower exposure in scenarios dominated by an increase of temperatures, and higher exposure for scenarios in which heavy precipitation events occur after pesticide application. Our case study shows that pesticides such as azoxystrobin, difenoconazole and MCPA are posing high ecological risks for aquatic organisms, and that the implementation of the ‘Farm-to-Fork’ strategy is crucial to reduce them, although will need additional measures to eliminate them.

## 1. Introduction

Mediterranean coastal wetlands have been considered biodiversity hotspots and play a key role in ecosystem service provision (Pérez-Ruzafa & Marcos, 2008). Several studies show that these ecosystems are impacted by a wide range of anthropogenic stressors (Martínez-Megías & Rico, 2022; Newton *et al*., 2014). Pollution by agricultural pesticides is one of the most important ones (Barbieri *et al*., 2020; Barhoumi *et al*., 2014; Calvo *et al*., 2021; Ccanccapa *et al*., 2016), however their impacts on aquatic ecosystem structure and biodiversity have been seldom investigated (Martínez-Megías & Rico, 2022). Some studies report that the presence of pesticides can have significant effects on aquatic communities (Pitacco *et al*., 2020), affecting food web stability and the relative abundance of predator and prey species (Quintana *et al*., 2018). Also, pesticide risks could significantly vary over time, becoming critical in some periods of the year due to interactions with other natural or anthropogenic stressors related to global climate change (Duchet *et al*., 2010).

The last report by the Intergovernmental Panel on Climate Change has predicted a temperature increase of up to 5.6 degrees for the Mediterranean region by the end of 21^st^ century, which is accompanied by a decrease in annual precipitations and an increasing occurrence of extreme events such as severe droughts and heatwaves (Ali *et al*., 2022). Although some authors have found that climate change could notably influence the environmental fate and toxicity of pesticides (Arenas-Sánchez *et al*., 2019; Holmstrup *et al*., 2010; Vilas-Boas *et al*., 2021), others indicate lower side-effects in water bodies due to increasing biodegradation (Willming & Maul, 2016). Thus, there is no apparent consensus on whether climate change is expected to increase or decrease pesticide risks for aquatic ecosystems.

The positive effect of warming on pesticide degradation rates could be compensated by an increasing occurrence of agricultural pests. In fact, several studies show that, under a climate change scenario, some agricultural pests could spread beyond their original distribution areas (Eitzinger *et al*., 2013), and become more prevalent in a higher number of agricultural crops (Noyes *et al*., 2009). Therefore, it is expected that many farmers tend to increase pesticide dosages per cropland area in regions with a significant increase in temperatures or precipitation (Delcour *et al*., 2015; Hader *et al*., 2022). This is supported by the fact that, despite legal restrictions have been put in place in many regions of Europe, this has not been translated into a reduction on the total amount of pesticides used in agriculture (Lamichhane *et al*., 2016).

On the other hand, the continued reliance on agricultural pesticides and their environmental impacts has been addressed through the enactment of the European Green Deal by the European Commission. This policy includes the ‘Farm-to-Fork’ strategy, which aims for a 50% reduction in pesticide usage by 2030 (European Commission, 2021). Therefore, it is expected that the future environmental risk of pesticides will be influenced by two key variables: climate change, which will affect pesticide exposure patterns, and pesticide management, which can result in an increase or decrease of pesticide dosages given the prevalence of agricultural pests or the implementation of environmental protection policies such as the ‘Farm-to-Fork’ strategy. The consequences of these two key variables for aquatic ecosystems have been scarcely investigated and need to be addressed making use of prospective risk models and realistic environmental scenarios.

During the last two decades, there has been notable progress in the use of mathematical models to assess pesticide exposure concentrations and ecological risks (e.g. EUFRAM, 2006; Rico *et al*., 2021; Maertens *et al*., 2022). Among them, Bayesian network approaches have arisen as innovative and flexible tools to explore the influence of different agricultural management scenarios on pesticide risks (Kaikkonen *et al*., 2021). Bayesian networks are probabilistic graphical models composed of nodes connected through arcs representing conditional probability tables (Aguilera *et al*., 2011; Kaikkonen *et al*., 2021). Based on new evidence, this direct acyclic graph uses the Bayes’ rule to update the probability distributions of the network nodes (Carriger *et al*., 2016; Kanes *et al*., 2017). Furthermore, they can function as meta-models, integrating information and knowledge from several sources and sub-models into a single predictive tool. Differently to traditional regulatory approaches, these probabilistic approaches enable better consideration of uncertainty, and can incorporate spatial and temporal variability into pesticide exposure distributions, as well as distributions that represent the sensitivity of the non-target species potentially affected by the pesticides (Mentzel *et al*., 2022; Piffady *et al*., 2021). Despite their potential to assess ecological risks given the influence of different pesticide management scenarios, their application to assess the influence of climate change or the implementation of new environmental policies is yet very limited (Kaikkonen *et al*., 2021; Moe *et al*., 2021).

This study aimed to assess how changes in future climate and pesticide use practices can influence the ecological risks of pesticides in a protected Mediterranean coastal wetland impacted by intensive rice farming. Our study serves as a basis to understand how future temperature and precipitation patterns, in combination with changes in pesticide dosages corresponding to different pest resistance scenarios and the implementation of the ‘Farm-to-Fork’ strategy, can affect pesticide exposure and risks for aquatic ecosystems. The assessment shown here is grounded on probabilistic risk assessment and allows calculation of acute and chronic ecological risk distributions using a Bayesian network approach. This work is one of the first assessing the risks of pesticide pollution from rice cultivation areas in the Mediterranean region at large spatial and temporal scales and provides recommendations as to which pesticides should be targeted in future monitoring programs.

## 2. Materials and methods

### Study area

The Albufera Natural Park (ANP) is located in eastern Spain (**Fig. 1**) and is one of the most studied Mediterranean coastal wetlands (Martínez-Megías & Rico, 2022). It is formed by a coastal lagoon surrounded by intensive rice production and a marsh area (Soria, 2006). The ANP joined the Natura 2000 as Site of Community Importance (SCI) in 1989 and is considered a Special Birds Protection Area (SBPA), being one of the most important Ramsar wetlands of Spain (Paredes Losada, 2020). Despite its regulation status, the hydric regime of the lagoon is forcedly determined by the stages of rice cultivation and the water quality shows a poor ecological status due to the intensive use and emission of fertilizers and pesticides (Vera-Herrera *et al*., 2021). Several types of herbicides are applied in order to control gramineous weeds (*e.g. Echinocloa sp., Leptochloa sp..*), Cyperaceae (*Chara sp.*) and wild rice. Insecticides are also used to combat different aphid species, while fungicides are intensively applied to combat the rice blast fungus (*Magnaporthe grisea*).

**Figure 1.**
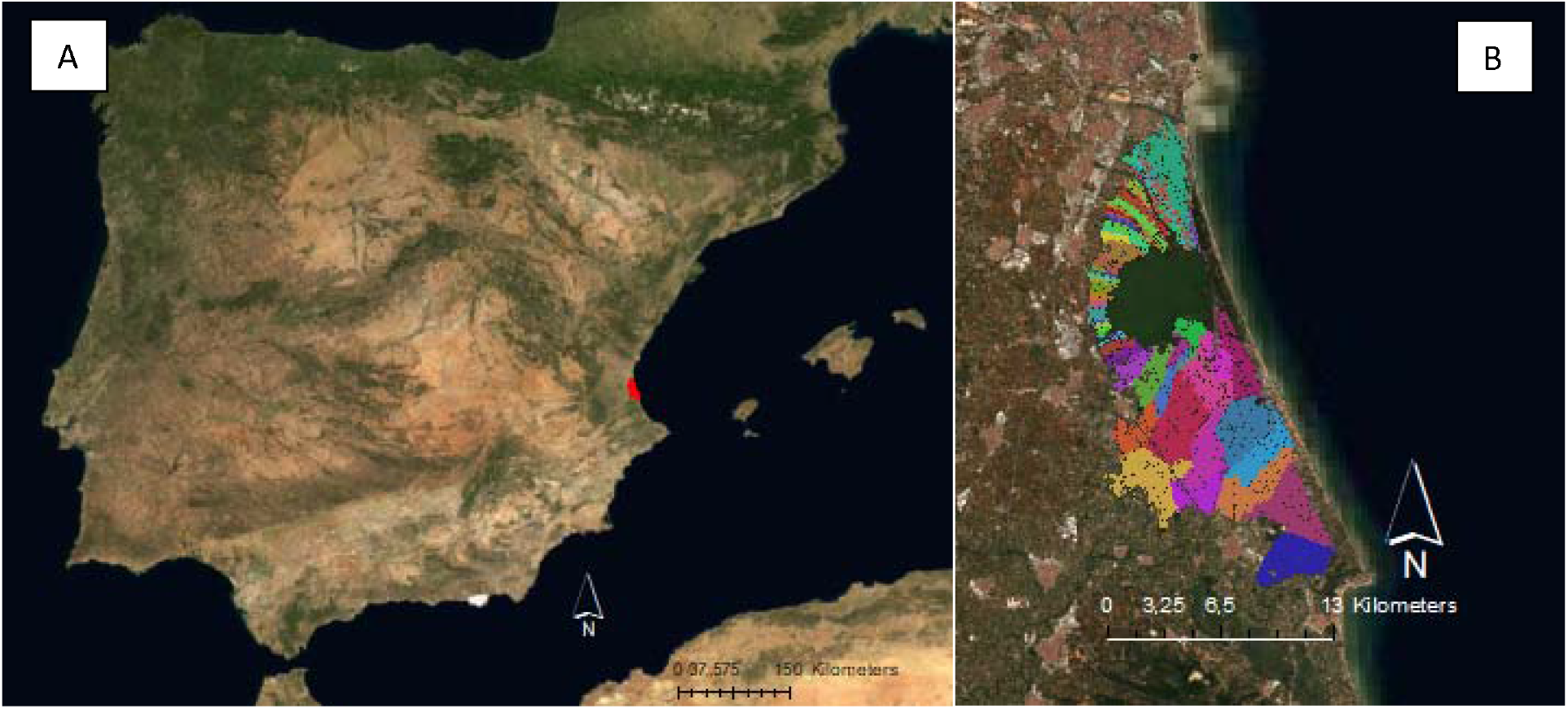
A: location of ANP in the Iberian Peninsula. B: View of the Albufera coastal lake and their surrounding rice plots forming the ANP. Colored polygons indicate hydrological clusters (i.e., rice plots with similar water renewal rate and water balance) used in the pesticide exposure modelling.

In the ANP, rice is planted between May and September, with narrow time variations depending on farmers’ management. Seeding is done with dried fields. Plant germination takes place one week after seeding. The water depth in the rice fields is maintained through constant irrigation at approximately 10 cm during the whole growing season, except for four emptying events, which are required for the application of the pesticides. The first three emptying events correspond with herbicide applications that take place between 1 and 6 weeks since rice seeding and have a duration of approximately three days. During early July (8 weeks since seeding), the water fields are emptied for approximately 7 days. The purpose of this last drainage process (locally called *eixugó*) is to prevent the proliferation of some competing plant species such as *Echinocloa sp*., as well as the application of herbicides and insecticides. Fungicides are applied in late summer by helicopter without water removal. Finally, rice harvest takes place around the last week of September, after the paddy fields have been completely dried. The rice cultivation phases, and pesticide applications, are summarized in **Fig. 2**.

**Figure 2.**
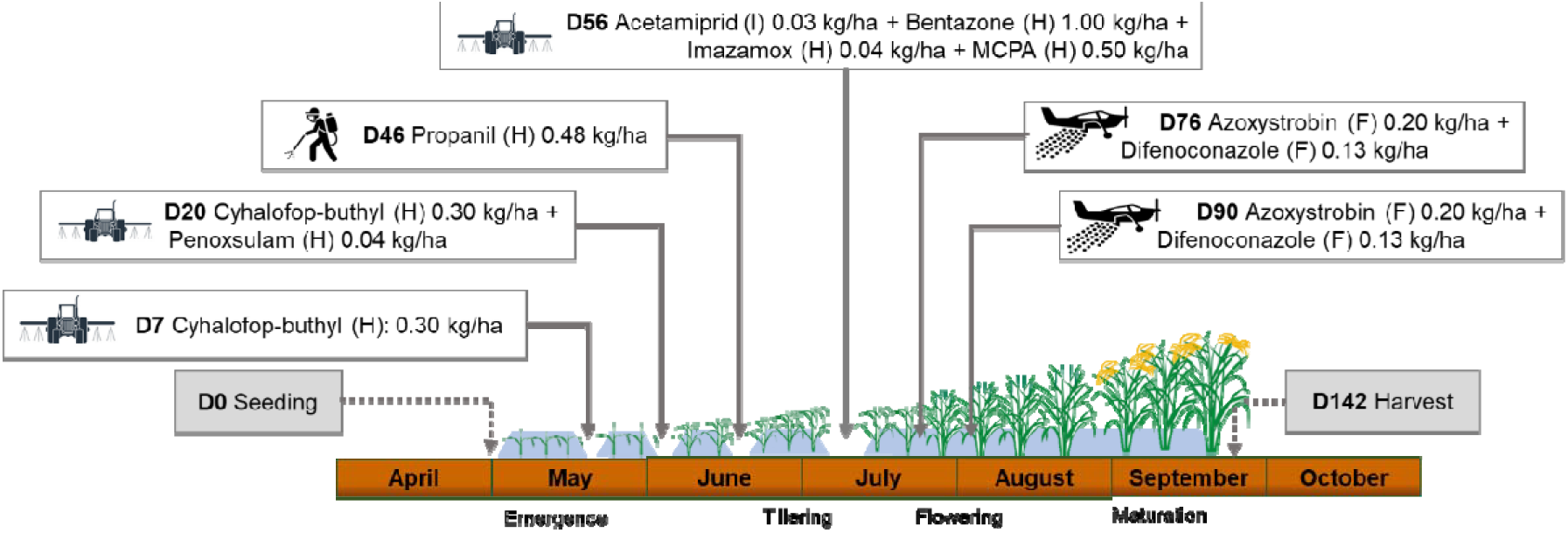
Rice cultivation phases and pesticide applications in the rice fields of the Albufera Natural Park, including mode of application (i.e., truck, back-pack, helicopter) and dosage. D: day relative to seeding; F: fungicide; H: herbicide; I: insecticide.

### Study design and scenarios

Pesticide risks were assessed for the nine pesticides applied during the rice growing season (**Fig. 2**) using nine scenarios that describe differences between current and future climate conditions and pesticide application (**Table 1**). Weather data representing each climatological scenario were obtained for the rice growing season (May-October) and consisted of mean daily temperature (°C), daily total precipitation (cm) and daily evapotranspiration (cm). Meteorological data for 2008 was obtained from ©AEMET (2021) and was used to build the ‘current’ baseline climate scenario, while 2050 and 2100 forecasts were used to predict pesticide exposure in paddies in the mid- and long-term, respectively. Weather projections were obtained from the Max Planck Institute Earth System Model at base resolution (MPI-ESM-LR, Giorgetta *et al*., 2013), as it is one of the few available models proposed by ©AEMET (2021) which has calculated weather forecasts for the closest meteorological station (Valencia, Spain). This model has been considered appropriate in other studies performed in the Júcar River Basin (Pool *et al*., 2021) and in other regions of Spain (Fernández *et al*., 2017). The model predictions are based on the Representative Concentration Pathway (RCP) 8.5 emission scenario, which represents the worst-case scenario for CO_2_ emissions through the 21^st^ century without considering any mitigation measures (Pool *et al*., 2021). The rest of emission scenarios were not included as they did not have precipitation data available. Pesticide dosages recommended by the manufacturers were used to simulate the ‘current’ (2008) and the 2050 and 2100 scenarios, while additional scenarios with increased pesticide and reduced pesticide dose were defined considering 50% more and less the recommended dose. The increased dosage is based on pesticide use trends indicated by local farmers, which claimed low pesticide efficacy to increasing pest prevalence and resistance, and the need to increase dosages to minimize the use of manpower for the elimination of unwanted weeds. The reduced dosage is based on the environmental target set by the European ‘Farm-to-Fork’ strategy to reduce agricultural dependance on pesticides and lower environmental risks (European Commission, 2021).

**Table 1.** Climate and pesticide use scenarios used in this study.

| Scenario name | Abbreviation | Input data |  |
| --- | --- | --- | --- |
|  |  | Meteorological data | Pesticide dose |
| Current | 2008 | 2008 | recommended |
| Mid-term projection | 2050 | 2050 | recommended |
| Long-term projection | 2100 | 2100 | recommended |
| Current, increasing pesticide dose | 2008+ | 2008 | 50% higher |
| Mid-term projection, increasing pesticide dose | 2050+ | 2050 | 50% higher |
| Long-term projection, increasing pesticide dose | 2100+ | 2100 | 50% higher |
| Current, reduced pesticide dose | 2008- | 2008 | 50% lower |
| Mid-term projection, reduced pesticide dose | 2050- | 2050 | 50% lower |
| Long-term projection, reduced pesticide dose | 2100- | 2100 | 50% lower |

### Pesticide exposure assessment

The rice production area of the ANP (210 km^2^) was divided into hydrological rice-production clusters. For this, the cadastral cartographic data of the study area was retrieved from the government of Spain (Spanish Government, 2021) and filtered to select only areas used for rice production. Based on cartographical data for the irrigation ditches of the Natural Park, we created polygons by segmenting the ditches longitudinally (segments of 500 m) and performing transversal buffering (from the center of the ditch, 500 m). Then, the centroid was calculated, and the rice paddies in the cadastral cartographic dataset were clustered based on the Euclidean distance to the nearest centroid of the segmented ditch, yielding a total of 552 rice production clusters (**Fig. 1**).

The data on the water irrigation rate of each rice production cluster was defined using a monitoring study of flow rates in 62 irrigation channels within the Natural Park in 2008 (Figure S1). Additionally, we set 8 points on the southern part of the Natural Park due to the paucity of measures in that area with the mean values of the sample. Renewal rates were assigned to the clusters based on its flow rate and total surface. The complete calculation of water renewal rates is shown in the Supplementary Material (Text S2).

The start date of the rice cultivation in each cluster was assigned following a stochastic approach. Ten cases were defined based on the starting day of the rice growing season (from May 3^rd^ to May 12^st^), maintaining constant the crop shift shown in **Fig. 2.** Then, based on the no spatial correlation with starting dates nor crop durations in the study area, those cases were assigned randomly to each of the 552 rice production clusters.

Pesticide exposure concentrations in each cluster was predicted using the RICEWQ model, version 1.92 (Waterborne Environmental Inc, 2022). Its core equations involve pesticide processes occurring in the air compartment (*i.e*., drift, volatilization, wash-off, crop interception), the aquatic compartment (*i.e*., biological and chemical decay, drainage, transformation, sorption) and the interphase between water and sediment (*i.e*., settling, resuspension, seepage, diffusion). The RICEWQ model runs were based on the study region’s specific rice-paddies, meteorological and hydrological parameters. The rice paddy parameters were obtained from field measurements and included some default values included in the RICEWQ manual (Tables S1 and S3). The meteorological data used for the RICEWQ simulations was the one described above for the selected scenarios. Since the selected prediction model did not have data for daily mean temperature, this parameter was estimated from the arithmetic mean of the daily maximum and minimum temperature. Furthermore, the daily mean temperature data obtained were used to calculate the evapotranspiration data. Details of evapotranspiration calculations are provided in Supplementary Material (Text S3). Water irrigation and drainage were set by the renewal rate assigned to each of the clusters, the gains (rainfall) and losses (evapotranspiration) of the selected scenarios to keep a water depth of 10 cm during the rice growing season. The water balance also considered the four drainage events (D7, D20, D46, D56) for pesticide application described in **Fig. 2.** The equations used to derive the water balance are provided in the Supplementary Material (Text S1). Physico-chemical data for the nine pesticides evaluated in this study were mainly obtained from the Draft Assessment Reports (DAR) published by the European Food Safety Authority (EFSA) and the Pesticide Properties Database (PPDB). These are shown in the Supplementary Material (Table S2).

Pesticide exposure concentrations in the paddy field water for each of the 9 pesticides were assessed for the 552 clusters in each of the 9 scenarios described in Table 1, yielding a total of 44712 model runs. To perform all these model runs, a handler for the RICEWQ, named autoRICEWQ, was developed. This software automatically creates the input files, executes RICEWQ and process the output data for each run. All the details for the autoRICEWQ generated for this study can be found at Fuentes-Edfuf & Martínez-Megías (2022) (open source under GPL-3.0 License, programmed in Python 3). From the predicted pesticide exposure concentrations in the paddy field, we calculated the Peak Exposure Concentration (PEC) and the Time Weighted Average Concentration over a period of 21 days (TWAC21).

### Pesticide effect assessment

Laboratory toxicity data for primary producers, invertebrates, and vertebrates were retrieved from the ECOTOX database (EPA, 2021). The available data were screened and classified according to the criteria described by Rico et al. (2019). Briefly, acute toxicity data for vertebrates were based on 2-4 days LC50 values, for invertebrates on 2-4 days LC50 or EC50 (immobilization), and for primary producers on EC50 (growth rate inhibition or yield) using an exposure period of 3-5 days for algae and more than 7 days for macrophytes. Chronic toxicity data for vertebrates were based on EC10 or NOECs (growth rate, development, behavior, mortality, immobilization) for an exposure period higher than 21 d, for invertebrates on EC10 or NOECs using similar endpoints and exposure durations as for vertebrates, and for primary producers based on EC10 or NOEC values for the same exposure duration and endpoints as for the acute assessment.

Acute and chronic Species Sensitivity Distributions (SSDs) were built using the available toxicity data by fitting a log-normal distribution with the MOSAIC software (King *et al*., 2013). In most cases, there were toxicity values for 8 or more taxa to build the acute or chronic SSDs; however, when there was toxicity data for less than 8 taxa, acute-to-chronic extrapolations were applied. In this way, acute SSDs were complemented with chronic EC10 or NOECs for unrepresented taxa by multiplying them by a factor of 100 for invertebrates and vertebrates, and 10 for primary producers. Chronic SSDs were complemented with acute EC50 or LC50 values by dividing them by a factor of 100 for invertebrates and vertebrates, and 10 for primary producers. The mean and standard deviation of the calculated SSDs for each pesticide are shown in Table S4.

### Probabilistic risk assessment

Risk Quotients (RQs) were predicted using Bayesian networks built with the Netica software (Norsys Software Corp., www.norsys.com) using the guidelines provided by Marcot *et al*., (2006) and Pollino & Henderson (2010). The Bayesian network followed a simplified structure of the one carried out by Mentzel et al. (2022), and was composed of eight nodes (**Fig. 3).** The exposure concentration distribution was determined by the scenario combination and the exposure time (acute or chronic). Model pesticide Exposure Concentration Distributions (ECDs) were fitted using the PEC or TWAC21 pesticide exposure concentrations obtained with the autoRICEWQ runs for the different scenarios. The fitting of the ECDs was performed with the *fitdistrplus* R package (Delignette-Muller & Dutang, 2015) for the model distributions available in Netica (Table S4), and the best fitting distribution was selected using the Akaike Information Criterion.

**Figure 3.**
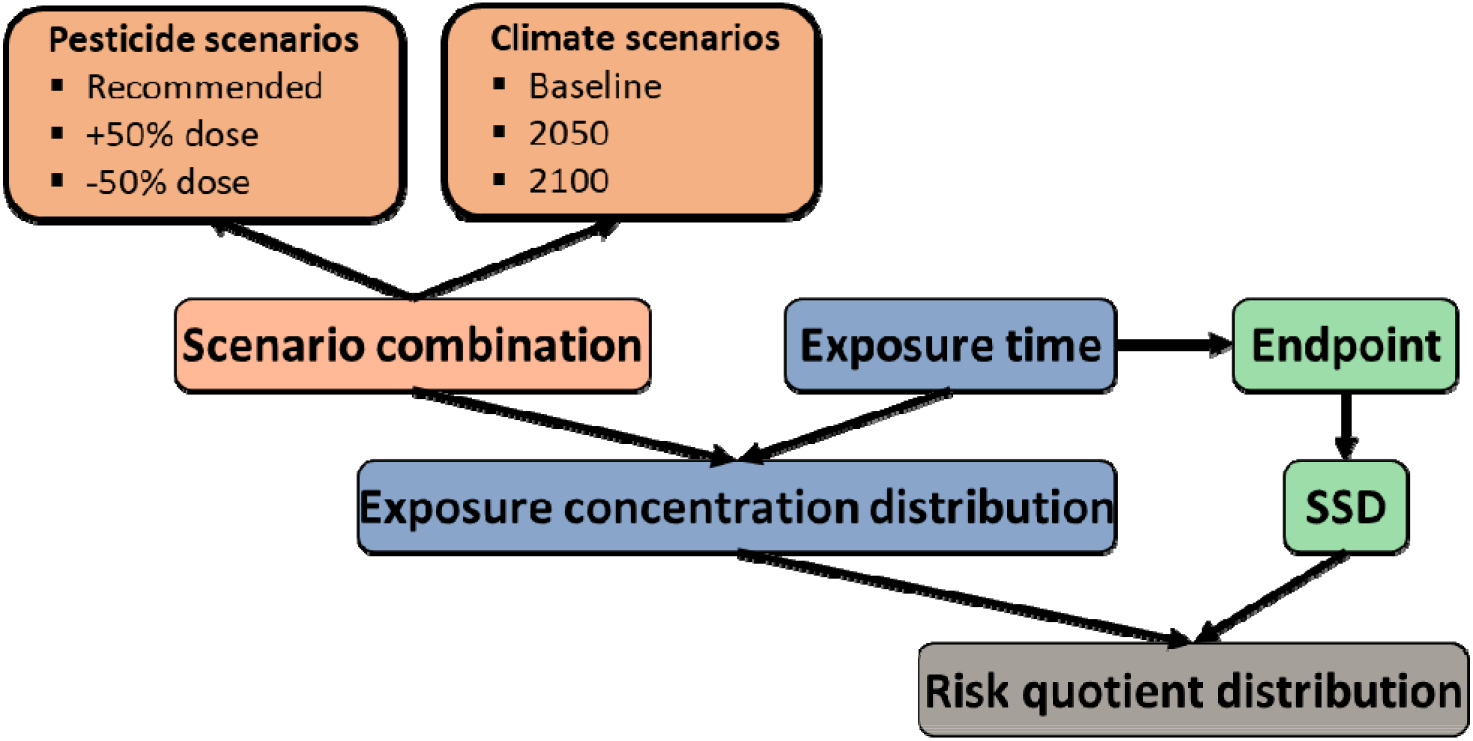
Conceptual model of the Bayesian Network (BN) approach used in this study. The full description of the BN nodes is shown in Table S5.

RQs distributions were calculated as the ratio between values within the ECD and the SSD for the pesticides, considering the PEC/SSD acute and the TWAC21/SSD chronic for the acute and chronic risk assessments, respectively. RQs below 0.1 were considered to result in no risks for aquatic ecosystems; RQs larger than 0.1 and lower than 1 were considered to result in potential risks; RQs between 1 and 10 were assumed to represent moderate risks, and RQs over than 10 were considered to pose high risks. The RQ distribution was visualized with a stock bar to better compare the percentage of cases (i.e., rice production clusters) assigned to each risk category in the different scenarios.

## 3. Results and discussion

### Pesticide exposure assessment

The three climate scenarios were significantly different, both in terms of daily mean temperature and total precipitation for the whole crop season (**Fig. 4).** The mean daily temperature (±SD) was 2O.9±4.1°C for 2008, 23.7±4.78°C for 2050 and 27.41±4.4°C for 2100. Regarding annual precipitation, it amounted to 1759 mm for 2008, 1301 mm for 2050, and 784 mm for 2100. The results of the exposure assessment show that the different weather projections for 2008, 2050 and 2100 notably affected pesticide exposure distributions (**Fig. 5).** For most pesticides, the increase in temperatures and the reduction in precipitation of the 2050 and 2100 scenarios resulted in a decrease of predicted exposure concentrations, which in general was more evident for the PEC distributions as compared to the TWAC ones. However, for other pesticides such as acetamiprid or imazamox, the exposure distributions in the different time horizons were rather comparable, while propanil showed higher PEC and TWAC distributions in 2050 as compared to 2008 and 2100 (**Fig. 5).**

**Figure 4.**
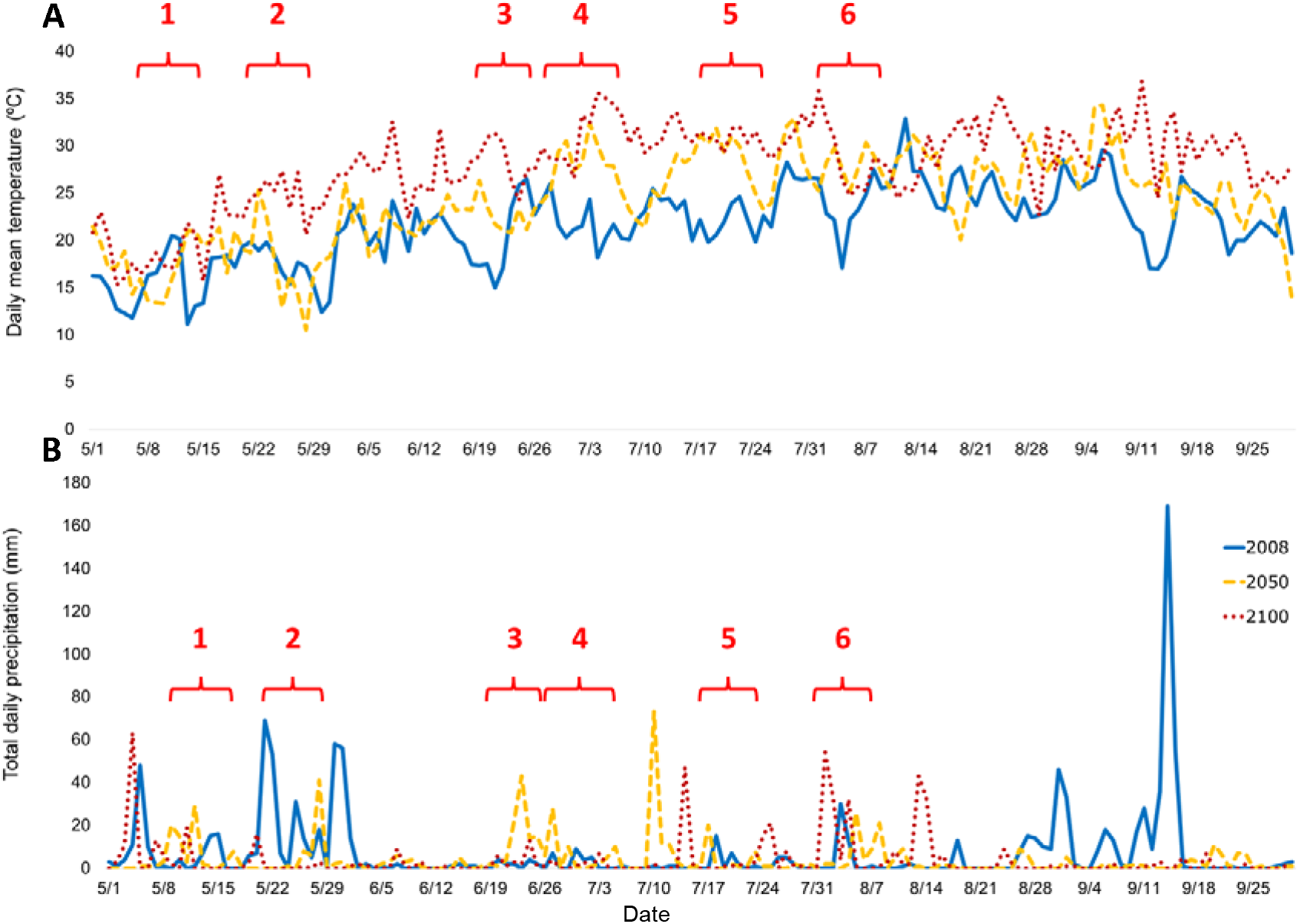
Variation of mean daily temperature (A) and total daily precipitation (B) over the rice cultivation period. Each line represents a different climate scenario (i.e., 2008, 2050, 2100) predicted by the MPI-ESM-LR model. The red numbers indicate pesticide application events: 1: cyhalofop (1^st^) (D7); 2: cyhalofop (2^nd^) and penoxsulam (D20); 3: propanil (D46); 4: acetamiprid, bentazon, imazamox and MCPA (D56); 5: azoxystrobin and difenoconazole (1^st^); 6: azoxystrobin and difenoconazole (2^nd^). See Fig. 1 for a detailed description of the pesticide dosages and modes of application.

**Figure 5.**
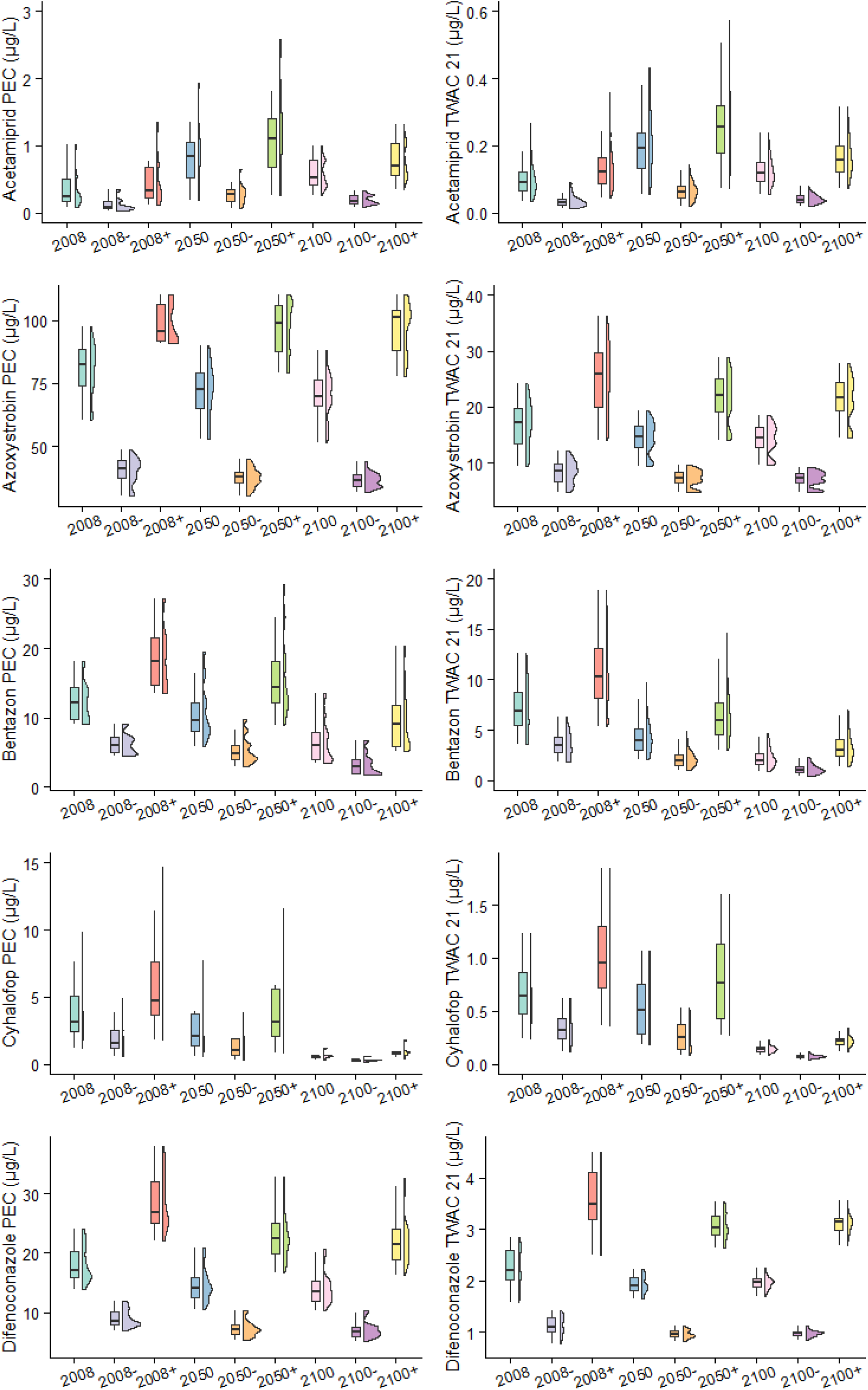
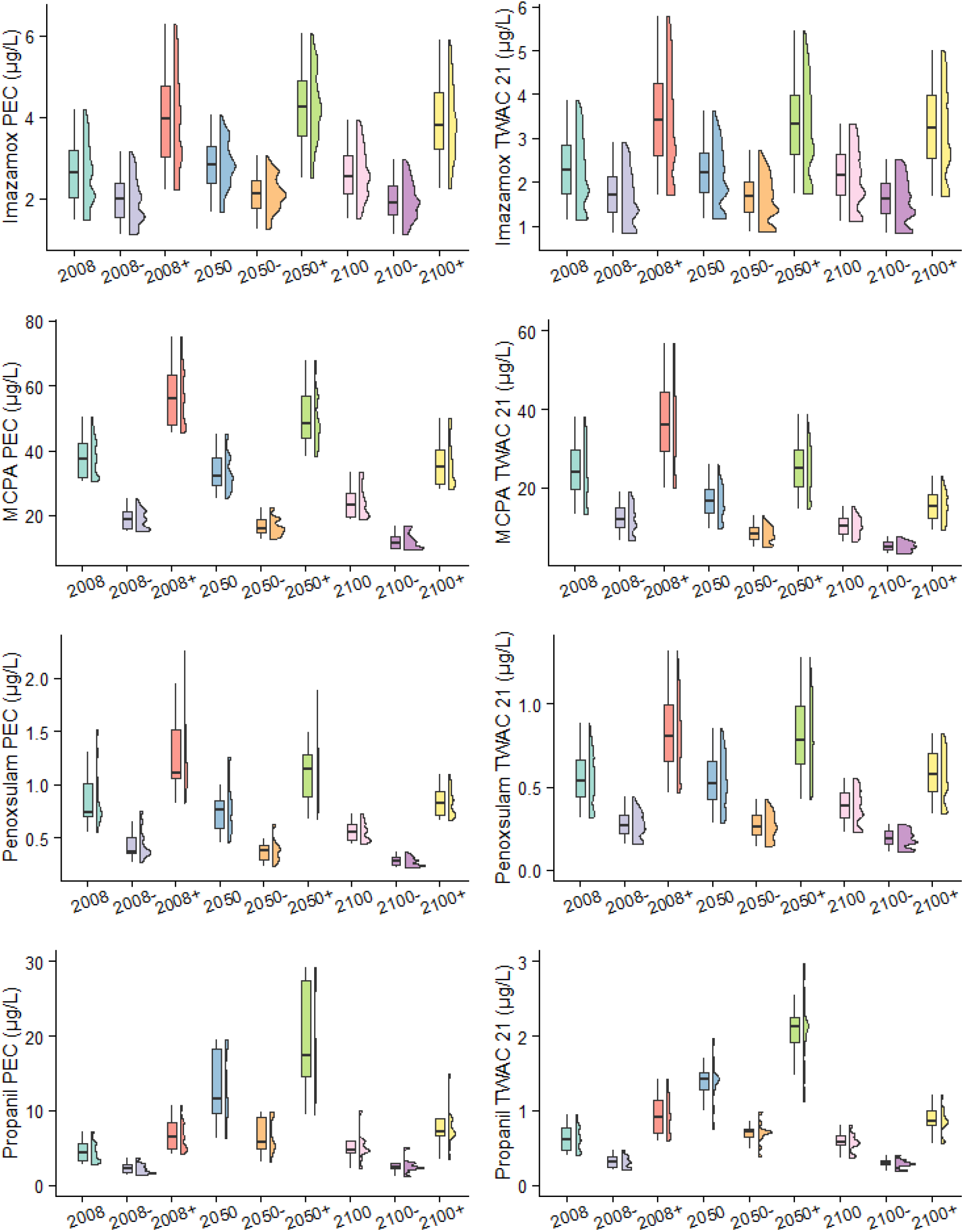
Peak Exposure Concentration (PEC) and highest Time Weighted Average Concentration (TWAC) distributions for the nine pesticides evaluated in this study within each scenario. The box plot shows the median of the distribution (black line) as well as the 25^th^ and 75^th^ percentiles (lower and upper boundaries, respectively). See **Table 1** for a description of the pesticide scenario abbreviations.

The observed trend towards a reduction of PEC and TWACs for most pesticides in the 2050 and 2100 scenarios could be related to processes such as volatilization (Bloomfield *et al*., 2006; Noyes *et al*., 2009) and microbial biodegradation in water or sediment (Delcour *et al*., 2015) These processes were enhanced by increasing temperatures as described in the equations that support the RICEWQ calculations. For compounds such as acetamiprid or imazamox, the slight differences in PEC or TWACs among the three environmental scenarios could be related to their application type and their specific physico-chemical properties. For example, imazamox is applied directly to dried soils and shows a very low degradation rate in soil, therefore temperature is expected to affect much less this process. Acetamiprid, is also applied directly to soil during the 7-day dry period (i.e., the *eixugó* period). For both pesticides, PECs seem to be driven by soil resuspension after rewetting, a process that is related to agricultural irrigation practices. As for propanil, the higher water concentration predicted for the 2050 scenario is closely related to heavy precipitation events occurring during or shortly after the pesticide application date (around D46). Other factors that may have affected pesticide exposure include the temporal decrease in daily mean temperature, a reduction in evapotranspiration, and an increase in sediment resuspension and pesticide wash-off from the rice plants. In line with this, some studies, such as Nolan *et al*. (2008) or Lewan *et al*. (2009) indicate the timing of precipitation events in relation to pesticide application dates and drainage losses as one of the main drivers of peak exposure concentrations in water bodies.

Regardless of the climatic projections, variations in pesticide application dosages resulted in marked PEC and TWAC distribution differences as compared to the recommended dosages (**Fig. 5).** Differences in exposure distributions resulting from the different dosage scenarios were particularly noticeable for PECs of the fungicides azoxystrobin and difenoconazole, which are applied in periods in which the paddy fields were filled with water, so that a fraction of the applied dosage is directly dissolved into the rice plot water. Interestingly, variations in pesticide contamination in paddy field water related to the different application scenarios tested in this study were more prominent than variations in pesticide exposure related to the 2050 and 2100 weather projections provided by the RCP 8.5 emission scenario. This suggests that, within this century, pesticide use management is likely more important than climate change factors to determine environmental exposure.

### Ecological risk assessment

The Bayesian network predicted acute and chronic RQ distributions as the ratio between the ECDs and the SSDs (**Fig. 6).** For most pesticides, the percentage of cases with moderate or high acute risks in the baseline scenario was relatively low (<5%). Exceptions were the fungicides azoxystrobin and difenoconazole, with a probability of 16% of the cases showing a moderate risk. Regarding chronic risks, for the baseline scenarios, azoxystrobin, difenoconazole, MCPA, imazamox and penoxsulam had a probability of 5% of the cases of being at a moderate to high risk level. The compounds posing the largest chronic risks were azoxystrobin, difenoconazole, and MCPA, with 54%, 21% and 13% of the ANP rice clusters showing moderate or high chronic risks, respectively (**Fig. 6).**

**Figure 6.**
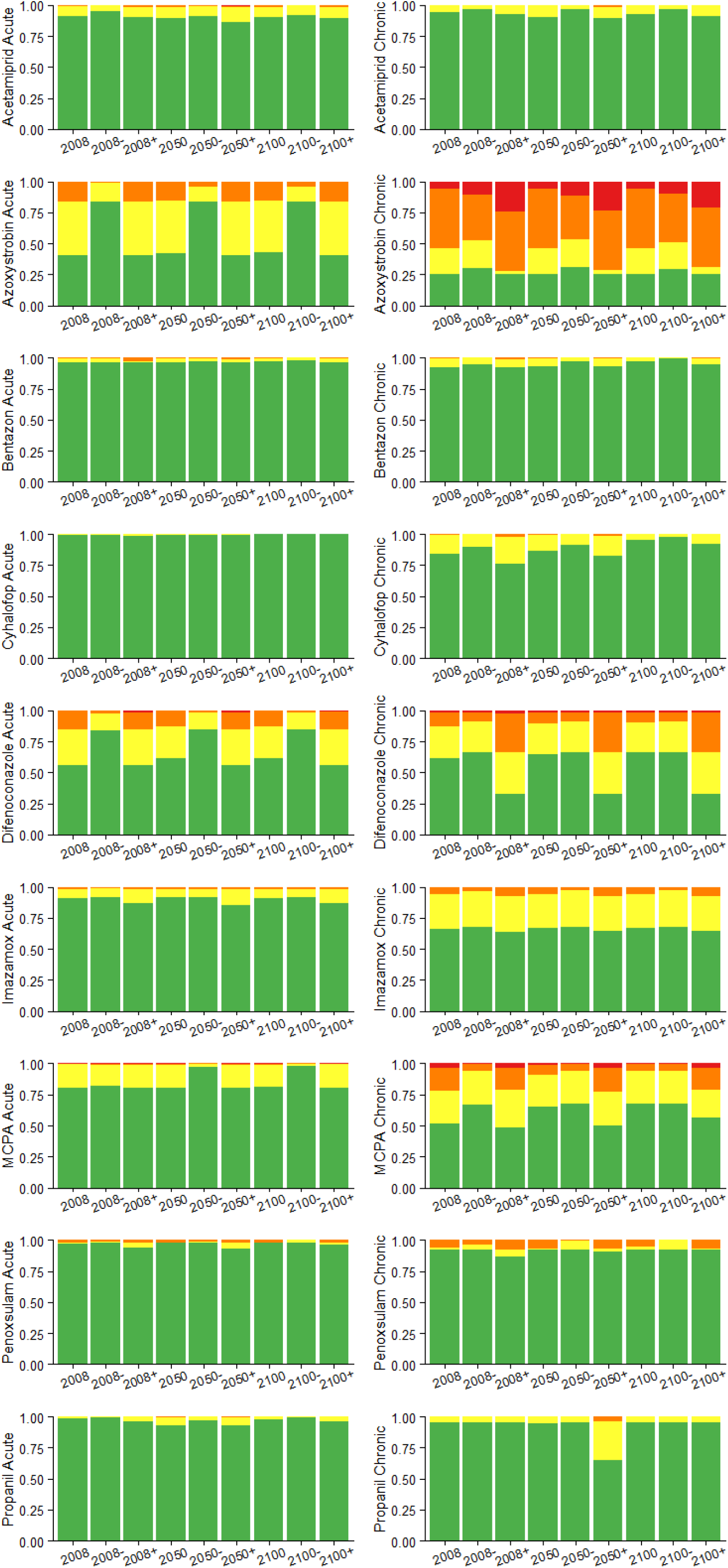
Bar plots showing the fraction of acute and chronic Risk Quotients (RQs) for the different rice production clusters falling within each risk category. The colors indicate the risk categories: Green: no risk (RQs: 0-0.1); Yellow: potential risk (RQs: 0.1-1); Orange: moderate risk (RQs: 1-10); Red: high risk (RQs: 10-10000). **See Table 1** for a description of the pesticide scenario abbreviations.

The different climate projections (2050 and 2100) had a mild influence on the RQ distributions, despite the decrease in exposure concentrations described for some compounds in the previous section. The most noticeable influence was observed for the chronic RQ distribution of the herbicide MCPA, with a probability of rice clusters showing moderate or high risks decreasing from 21% in the 2008 scenario, to 9% in the 2050 scenario, and 6% in the 2100 scenario. For propanil, the acute RQ distribution changed according to the PEC increase in 2050, indicating a larger probability of potential risks (7% of cases) in that scenario as compared to the 2008 (1%) and the 2100 (2%) ones (**Fig. 6).**

The different pesticide dosage scenarios had a clear influence on the RQ distributions for the pesticides, particularly for those showing moderate and high risks in the 2008 scenario. For example, the percentage of rice production clusters showing moderate or high chronic risks for azoxystrobin in the scenario accounting for a reduction of 50% of the dosage in the baseline scenario (i.e., 2008-) were 48%, while the percentage in the scenario simulating a 50% increase of the dose (i.e., 2008+) was 72%. In line with the mild influence of the weather scenarios described above, the temporal changes in the RQ distribution of scenarios assuming a 50% increase or decrease in the dosages was relatively low (**Fig. 6**).

### Implications for risk assessment and way forward

This study shows how weather projections and environmental management strategies can be integrated into a probabilistic framework to assess current and future risks of pesticides in a Mediterranean wetland of high ecological value. The approach allows integration of spatial-temporal variability in terms of hydrological regimes, weather conditions, and pesticide application schemes, and complements environmental monitoring studies that have shown unacceptable pesticide risks for aquatic organisms in the same study area (Calvo *et al*., 2021).

Our results show that the fungicides azoxystrobin and difenoconazole, and the herbicide MCPA pose the largest ecotoxicological risks. Azoxystrobin and difenoconazole were introduced in the ANP as replacements of more toxic (prochloraz, tebuconazole), or recently banned compounds (carbendazim) (Andreu Sánchez, 2008). However, as shown in this study, short and long-term ecological risks in the rice production area of the ANP may be expected. Semi-field experiments performed in Sweden and the Netherlands show chronic toxic effects of azoxystrobin at concentrations that are an order of magnitude lower than the TWACs calculated in this study, with copepods and some Cladocera showing the largest abundance declines (Gustafsson et al., 2010; van Wijngaarden et al., 2014). Difenoconazole has proven to be very toxic to daphnids (Moreira *et al*., 2020), fish and algae (Man *et al*., 2021) in other ecosystems impacted by rice production (Shen *et al*., 2022). On the other hand, MCPA is relatively toxic to eelgrass and dicotyledonous aquatic plants (Nielsen & Dahllöf, 2007), and has been highlighted as one of the most toxic compounds in other rice production areas such as the Ebro Delta in Spain (Barbieri *et al*., 2020).

Our study has demonstrated that climate conditions can influence pesticide exposure and risks. In this case-study, local precipitation events occurring around the time of pesticide applications were found to be more determinant than temperature increases forecasted for the next decades. Therefore, further attention should be paid to integrate changes in precipitation regimes, including extreme rainfall events, into future pesticide risk assessment scenarios for the Mediterranean region.

The outcomes of the risk assessment show that the implementation of environmental protection measures, such as the dose reduction measure promoted by the ‘Farm-to-Fork’ strategy, will be key to reduce the aquatic risks of pesticides in the next decades. However, the reduction in 50% of the dose does not completely exclude risks for some pesticides. Additional risk reduction measures, such as the replacement of highly hazardous substances, the incorporation of integrated pest management practices or the construction of constructed wetlands can limit pesticide emissions to surrounding water bodies (Alexoaei et al., 2022; Martín et al., 2020; Pavlidis & Karasali, 2020; Silva et al., 2022). A recent study by Rodrigo et al. (2022) shows that two constructed wetlands located next to rice production areas can reduce metal loads and the number of pesticides entering the Albufera Lake. Further studies should be developed to calculate pesticide transport in the drainage ditches of the ANP and potential risks for aquatic communities in the Albufera Lake under different weather and pesticide management scenarios.

The modelling approach described here offers opportunities to predict pesticide risks in a complex spatial-temporal environmental setting; however, it has some caveats. On the one hand, it disregards the formation of hazardous pesticide metabolites and transformation products. Therefore, the higher dissipation of the parent compounds caused by the forecasted increase in temperatures results in lower exposure levels and risks. Although the RICEWQ model includes some processes to account for this, we found that the amount of chemical fate and toxicity data to characterize exposure and risks of key pesticide metabolites was too limited. Therefore, this aspect could not be included as part of this study.

Another major limitation is that we considered the sensitivity of aquatic ecosystems, represented by the acute and chronic SSDs constant over the 2050 and the 2100 climate scenarios. Some studies suggest that aquatic organisms will have a higher sensitivity to pesticide exposure in scenarios of elevated (+5°C) temperature (Camp & Buchwalter, 2016; Roth et al., 2022; Vilas-Boas et al., 2021). Therefore, our risk projections may have, to some extent, underestimated the actual ecological risks. On the other hand, climate change, and the extreme weather events associated to it are expected to affect the structure of aquatic communities (Polazzo et al., 2022), filtering for species assemblages that may be more (or less) sensitive to different pesticides. These aspects should be further investigated and potentially incorporated into future pesticide risk projections for the Mediterranean region.

## 4. Conclusions

This study shows how Bayesian network approaches can be used to assess the influence of different climate change and pesticide management scenarios on the ecological risks of pesticides. The case-study performed here for the nine pesticides used in rice production in the ANP shows that future climate projections will result in lower exposure and risk distributions in scenarios dominated by an increase of temperatures, while exposure and risks can increase for some pesticides applied during periods of heavy precipitation events, which will be more recurrent in the future. Moreover, it shows that three out of the nine evaluated pesticides (azoxystrobin, difenoconazole and MCPA) pose high ecological risks for aquatic organisms and should be included in further ecotoxicological experiments and monitoring programs in the study area. Finally, we have demonstrated that the increase of pesticide dosages due to the highest prevalence of agricultural pests is likely to have profound implications for aquatic biodiversity in coastal Mediterranean wetlands, and that the implementation of pesticide dose reduction programs, such as the European ‘Farm-to-Fork’ strategy, are crucial to reduce pesticide risks, although will need additional measures to completely eliminate them.

## Acknowledgments

This study was funded by the CICLIC project (Smart tools and technologies to assess the environmental fate and risks of Contaminants under Climate Change), funded by the Spanish Ministry of Science, Innovation and Universities (RTI 2018_097158_A_C32); the H2020-MSCA-ITN ECORISK2050 project, funded by the European Union’s Horizon 2020 research and innovation program (grant agreement No. 813124); and the Talented Researcher Support Programme - PlanGenT (CIDEGENT/2020/043) of the Generalitat Valenciana. We would like to thank J. Soria for the 2008 water flow data, and L. Blanch for the information on rice production and pesticide management practices.

## References

Agencia Estatal de Meteorología (©AEMET). Servicios climáticos. Análisis estacional València. (2021). Consulted on 02 December 2021. www.aemet.es.

Aguilera, P. A., Fernández, A., Fernández, R., Rumí, R., & Salmerón, A. (2011). Bayesian networks in environmental modelling. Environmental Modelling and Software, 26(12), 1376–1388. https://doi.org/10.1016/j.envsoft.2011.06.004.

Alexoaei, A. P., Robu, R. G., Cojanu, V., Miron, D., & Holobiuc, A. M. (2022). Good Practices in Reforming the Common Agricultural Policy To Support the European Green Deal – a Perspective on the Consumption of Pesticides and Fertilizers. Amfiteatru Economic, 24(60), 525–545. https://doi.org/10.24818/EA/2022/60/525.

Ali, E., Cramer, W., Carnicer, J., Georgopoulou, E., Hilmi, N., Le Cozannet, G., & Lionello, P. (2022). CCP 4 Mediterranean Region. In H.-O. Pörtner, D. C. Roberts, M. Tignor, E. S. Poloczanska, K. Mintenbeck, A. Alegría, M. Craig, S. Langsdorf, S. Löschke, V. Möller, A. Okem, & B. Rama (Eds.), Climate Change 2022: Impacts, Adaptation and Vulnerability. Contribution of Working Group II to the Sixth Assessment Report of the Intergovernmental Panel on Climate Change (pp. 2233–2272). Cambridge University Press. https://doi.org/10.1017/9781009325844.021.2233.

Andreu Sánchez, D. O. (2008). Evaluación de riesgos ambientales del uso de plaguicidas empleados en el cultivo del arroz en el Parque Natural de La Albufera de Valencia. Universitat Politècnica de València.

Arenas-Sánchez, A., López-Heras, I., Nozal, L., Vighi, M., & Rico, A. (2019). Effects of increased temperature, drought, and an insecticide on freshwater zooplankton communities. Environmental Toxicology and Chemistry, 38(2), 396–411. https://doi.org/10.1002/etc.4304.

Barbieri, M. V., Peris, A., Postigo, C., Moya-Garcés, A., Monllor-Alcaraz, L. S., Rambla-Alegre, M., Eljarrat, E., & López de Alda, M. (2020). Evaluation of the occurrence and fate of pesticides in a typical Mediterranean delta ecosystem (Ebro River Delta) and risk assessment for aquatic organisms. Environmental Pollution, 274. https://doi.org/10.1016/j.envpol.2020.115813.

Barhoumi, B., Clérandeau, C., Gourves, P. Y., Le Menach, K., Megdiche, Y. El, Peluhet, L., Budzinski, H., Baudrimont, M., Driss, M. R., & Cachot, J. (2014). Pollution biomonitoring in the Bizerte lagoon (Tunisia), using combined chemical and biomarker analyses in grass goby, Zosterisessor ophiocephalus (Teleostei, Gobiidae). Marine Environmental Research, 101(1), 184–195. https://doi.org/10.1016/j.marenvres.2014.07.002.

Bloomfield, J. P., Williams, R. J., Gooddy, D. C., Cape, J. N., & Guha, P. (2006). Impacts of climate change on the fate and behaviour of pesticides in surface and groundwater-a UK perspective. Science of the Total Environment, 369(1-3), 163–177. https://doi.org/10.1016/j.scitotenv.2006.05.019.

Calvo, S., Romo, S., Soria, J., & Picó, Y. (2021). Pesticide contamination in water and sediment of the aquatic systems of The Natural Park of the Albufera of Valencia (Spain) during the rice cultivation period. Science of The Total Environment, 774, 145009. https://doi.org/10.1016/j.scitotenv.2021.145009.

Camp, A. A., & Buchwalter, D. B. (2016). Can’t take the heat: Temperature-enhanced toxicity in the mayfly Isonychia bicolor exposed to the neonicotinoid insecticide imidacloprid. Aquatic Toxicology, 178, 49–57. https://doi.org/10.1016/j.aquatox.2016.07.011.

Carriger, J. F., Martin, T. M., & Barron, M. G. (2016). A Bayesian network model for predicting aquatic toxicity mode of action using two dimensional theoretical molecular descriptors. Aquatic Toxicology, 180, 11–24. https://doi.org/10.1016/j.aquatox.2016.09.006.

Ccanccapa, A., Masiá, A., Navarro-Ortega, A., Picó, Y., & Barceló, D. (2016). Pesticides in the Ebro River basin: Occurrence and risk assessment. Environmental Pollution, 211, 414–424. https://doi.org/10.1016/j.envpol.2015.12.059.

Delignette-Muller, M. L., & Dutang, C. (2015). fitdistrplus: An R package for fitting distributions. Journal of Statistical Software, 64(4), 1–34. https://doi.org/10.18637/jss.v064.i04.

Duchet, C., Caquet, T., Franquet, E., Lagneau, C., & Lagadic, L. (2010). Influence of environmental factors on the response of a natural population of Daphnia magna (Crustacea: Cladocera) to spinosad and Bacillus thuringiensis israelensis in Mediterranean coastal wetlands. *Environmental Pollution*, 158(5), 1825–1833. https://doi.org/10.1016/j.envpol.2009.11.008.

Eitzinger, J., Trnka, M., Semerádová, D., Thaler, S., Svobodová, E., Hlavinka, P., Šiška, B., Takáč, J., Malatinská, L., Nováková, M., Dubrovský, M., & Žalud, Z. (2013). Regional climate change impacts on agricultural crop production in Central and Eastern Europe - Hotspots, regional differences and common trends. Journal of Agricultural Science, 151(6), 787–812. https://doi.org/10.1017/S0021859612000767.

EUFRAM. (2006). Concerted action to develop a European Framework for probabilistic risk assessment of the environmental impacts of pesticides. papers3://publication/uuid/175535F4-D5D5-4E31-A1C7-2C09251FCF76

European Commission. (2021). The European Green Deal. http://eurlex.europa.eu/resource.html?uri=cellar:208111e4-414e-4da5-94c1-852f1c74f351.0004.02/DOC_1&format=PDF.

Fernández, J., Casanueva, A., Montávez, J. P., Gaertner, M. Á., Casado, M. J., Manzanas, R., & Gutiérrez, J. M. (2017). Regional climate projections over Spain: Atmosphere. Future Climate projections. CLIVAR Exchanges, 73(September), 39–44.

Fuentes-Edfuf, Y., & Martínez-Megías, C. (2022). backmind/autoRICEWQ: v1.0.2. https://doi.org/10.5281/ZENODO.5940235.

Giorgetta, M. A., Jungclaus, J., Reick, C. H., Legutke, S., Bader, J., Böttinger, M., Brovkin, V., Crueger, T., Esch, M., Fieg, K., Glushak, K., Gayler, V., Haak, H., Hollweg, H.-D., Ilyina, T., Kinne, S., Kornblueh, L., Matei, D., Mauritsen, T.,… Stevens, B. (2013). Climate and carbon cycle changes from 1850 to 2100 in MPI-ESM simulations for the Coupled Model Intercomparison Project phase 5. Journal of Advances in Modeling Earth Systems, 5(3), 572–597. https://doi.org/10.1002/jame.20038.

Gustafsson, K., Blidberg, E., Elfgren, I. K., Hellström, A., Kylin, H., & Gorokhova, E. (2010). Direct and indirect effects of the fungicide azoxystrobin in outdoor brackish water microcosms. Ecotoxicology, 19(2), 431–444. https://doi.org/10.1007/s10646-009-0428-9.

Hader, J. D., Lane, T., Boxall, A. B. A., Macleod, M., & Di, A. (2022). Science of the Total Environment Enabling forecasts of environmental exposure to chemicals in European agriculture under global change. Science of the Total Environment, 840(March), 156478. https://doi.org/10.1016/j.scitotenv.2022.156478

Holmstrup, M., Bindesbøl, A. M., Oostingh, G. J., Duschl, A., Scheil, V., Köhler, H. R., Loureiro, S., Soares, A. M. V. M., Ferreira, A. L. G., Kienle, C., Gerhardt, A., Laskowski, R., Kramarz, P. E., Bayley, M., Svendsen, C., & Spurgeon, D. J. (2010). Interactions between effects of environmental chemicals and natural stressors: A review. Science of the Total Environment, 408(18), 3746–3762. https://doi.org/10.1016/j.scitotenv.2009.10.067.

Kaikkonen, L., Parviainen, T., Rahikainen, M., Uusitalo, L., & Lehikoinen, A. (2021). Bayesian Networks in Environmental Risk Assessment: A Review. Integrated Environmental Assessment and Management, 17(1), 62–78. https://doi.org/10.1002/ieam.4332.

Kanes, R., Ramirez Marengo, M. C., Abdel-Moati, H., Cranefield, J., & Véchot, L. (2017). Developing a framework for dynamic risk assessment using Bayesian networks and reliability data. Journal of Loss Prevention in the Process Industries, 50, 142–153. https://doi.org/10.1016/j.jlp.2017.09.011.

King, G. K. K., Veber, P., Charles, S., & Delignette-Muller, M. L. (2013). MOSAIC _ SSD⍰: a new web-tool for the Species Sensitivity Distribution, allowing to include censored data by maximum likelihood. ArXiv Preprint ArXiv:1311.5772, 1–26.

Lamichhane, J. R., Dachbrodt-Saaydeh, S., Kudsk, P., & Messéan, A. (2016). Toward a Reduced Reliance on Conventional Pesticides in European Agriculture. Plant Disease, 100(1), 10–24. https://apsjournals.apsnet.org/doi/pdf/10.1094/PDIS-05-15-0574-FE.

Lewan, E., Kreuger, J., & Jarvis, N. (2009). Implications of precipitation patterns and antecedent soil water content for leaching of pesticides from arable land. Agricultural Water Management, 96(11), 1633–1640. https://doi.org/10.1016/j.agwat.2009.06.006.

Maertens, A., Golden, E., Luechtefeld, T.H., Hoffmann, S., Tsaioun, K. and Hartung, T. (2022). Probabilistic risk assessment–the keystone for the future of toxicology. Altex, 39(1), p.3.

Man, Y., Stenrød, M., Wu, C., Almvik, M., Holten, R., Clarke, J. L., Yuan, S., Wu, X., Xu, J., Dong, F., Zheng, Y., & Liu, X. (2021). Degradation of difenoconazole in water and soil: Kinetics, degradation pathways, transformation products identification and ecotoxicity assessment. Journal of Hazardous Materials, 418(June). https://doi.org/10.1016/j.jhazmat.2021.126303.

Marcot, B. G., Steventon, J. D., Sutherland, G. D., & Mccann, R. K. (2006). Guidelines for developing and updating Bayesian belief networks applied to ecological modeling and conservation. Canadian Journal of Forest Research, 36(12), 3063–3074.

Martín, M., Hernández-Crespo, C., Andrés-Doménech, I., & Benedito-Durá, V. (2020). Fifty years of eutrophication in the Albufera lake (Valencia, Spain): Causes, evolution and remediation strategies. Ecological Engineering, 155(April), 105932. https://doi.org/10.1016/j.ecoleng.2020.105932.

Martínez-Megías, C., & Rico, A. (2022). Biodiversity impacts by multiple anthropogenic stressors in Mediterranean coastal wetlands. Science of the Total Environment, 818, 151712. https://doi.org/10.1016/j.scitotenv.2021.151712.

Mentzel, S., Grung, M., Holten, R., Tollefsen, K. E., Stenrød, M., & Moe, S. J. (2022). Probabilistic risk assessment of pesticides under future agricultural and climate scenarios using a bayesian network. Frontiers in Environmental Science, 10(September), 1–17. https://doi.org/10.3389/fenvs.2022.957926.

Moe, S. J., Carriger, J. F., & Glendell, M. (2021). Increased Use of Bayesian Network Models has Improved Environmental Risk Assessments. Integrated Environmental Assessment and Management, 17(1), 53–61. https://doi.org/10.1002/ieam.4369.

Moreira, R. A., de Araujo, G. S., Silva, A. R. R. G., Daam, M. A., Rocha, O., Soares, A. M. V. M., & Loureiro, S. (2020). Effects of abamectin-based and difenoconazole-based formulations and their mixtures in Daphnia magna: a multiple endpoint approach. Ecotoxicology, 29(9), 1486–1499. https://doi.org/10.1007/s10646-020-02218-z.

Newton, A., Icely, J., Cristina, S., Brito, A., Cardoso, A. C., Colijn, F., Riva, S. D., Gertz, F., Hansen, J. W., Holmer, M., Ivanova, K., Leppäkoski, E., Canu, D. M., Mocenni, C., Mudge, S., Murray, N., Pejrup, M., Razinkovas, A., Reizopoulou, S.,… Zaldívar, J. M. (2014). An overview of ecological status, vulnerability and future perspectives of European large shallow, semi-enclosed coastal systems, lagoons and transitional waters. Estuarine, Coastal and Shelf Science, 140, 95–122. https://doi.org/10.1016/j.ecss.2013.05.023

Nielsen, L. W., & Dahllöf, I. (2007). Direct and indirect effects of the herbicides Glyphosate, Bentazone and MCPA on eelgrass (Zostera marina). Aquatic Toxicology, 82(1), 47–54. https://doi.org/10.1016/j.aquatox.2007.01.004.

Nolan, B. T., Dubus, I. G., Surdyk, N., Fowler, H. J., Burton, A., Hollis, J. M., Reichenberger, S., & Jarvis, N. (2008). Identification of key climatic factors regulating the transport of pesticides in leaching and to tile drains. Pest Management Science, 64, 933–944. https://doi.org/10.1002/ps.1587.

Noyes, P. D., McElwee, M. K., Miller, H. D., Clark, B. W., Van Tiem, L. A., Walcott, K. C., Erwin, K. N., & Levin, E. D. (2009). The toxicology of climate change: Environmental contaminants in a warming world. Environment International, 35(6), 971–986. https://doi.org/10.1016/j.envint.2009.02.006.

Paredes Losada, I. (2020). Presiones antrópicas y eutrofización en la marisma de Doñana y sus cuencas vertientes. Universidad de Sevilla.

Pavlidis, G., & Karasali, H. (2020). Natural Remediation Techniques for Water Quality Protection and Restoration. 327–340. https://doi.org/10.1007/978-3-030-48985-4_15.

Pérez-Ruzafa, A., & Marcos, C. (2008). Coastal Lagoons in the Context of Water Management in Spain and Europe. Sustainable Use and Development of Watersheds, 299–321. https://doi.org/10.1007/978-1-4020-8558-1_18.

Piffady, J., Carluer, N., Gouy, V., le Henaff, G., Tormos, T., Bougon, N., Adoir, E., & Mellac, K. (2021). ARPEGES: A Bayesian Belief Network to Assess the Risk of Pesticide Contamination for the River Network of France. Integrated Environmental Assessment and Management, 17(1), 188–201. https://doi.org/10.1002/ieam.4343.

Pitacco, V., Mistri, M., Ferrari, C. R., Sfriso, A., Sfriso, A. A., & Munari, C. (2020). Multiannual trend of micro-pollutants in sediments and benthic community response in a mediterranean lagoon (Sacca di Goro, Italy). Water (Switzerland), 12(4). https://doi.org/10.3390/W12041074.

Polazzo, F., Roth, S. K., Hermann, M., Mangold-Döring, A., Rico, A., Sobek, A., Van den Brink, P. J., & Jackson, M. C. (2022). Combined effects of heatwaves and micropollutants on freshwater ecosystems: Towards an integrated assessment of extreme events in multiple stressors research. Global Change Biology, 28(4), 1248–1267. https://doi.org/10.llll/gcb.15971.

Pollino, C. A., & Henderson, C. (2010). Bayesian networks⍰: A guide for their application in natural resource management and policy. Landscape Logic, Technical Report, 14, 48.

Pool, S., Francés, F., Garcia-Prats, A., Pulido-Velazquez, M., Sanchis-Ibor, C., Schirmer, M., Yang, H., & Jiménez-Martínez, J. (2021). From Flood to Drip Irrigation Under Climate Change: Impacts on Evapotranspiration and Groundwater Recharge in the Mediterranean Region of Valencia (Spain). Earth’s Future, 9(5), 1–20. https://doi.org/10.1029/2020EF001859.

Quintana, X. D., Boix, D., Gascón, S., & Sala, J. (2018). Management and restoration of mediterranean coastal lagoons in Europe (Càtedra d’). Gràfiques Agustí Printed.

Rico, A., Brock, T.C. and Daam, M.A. (2019). Is the effect assessment approach for fungicides as laid down in the European Food Safety Authority Aquatic Guidance Document sufficiently protective for freshwater ecosystems?. Environmental Toxicology and Chemistry, 38(10), pp.2279–2293.

Rico, A., Dafouz, R., Vighi, M., Rodríguez-Gil, J. L., & Daam, M. A. (2021). Use of Postregistration Monitoring Data to Evaluate the Ecotoxicological Risks of Pesticides to Surface Waters: A Case Study with Chlorpyrifos in the Iberian Peninsula. Environmental Toxicology and Chemistry, 40(2), 500–512. https://doi.org/10.1002/etc.4927.

Rodrigo, M. A., Puche, E., Armenta, S., & Juan, F. (2022). Two constructed wetlands within a Mediterranean natural park immersed in an agrolandscape reduce most heavy metal water concentrations and dampen the majority of pesticide presence. Available at: a5dea825-6b0a-42d2-bc55-8631b07a2bfd.pdf (researchsquare.com)

Roth, S. K., Polazzo, F., García-Astillero Honrado, A., Cherta, L., Sobek, A., & Rico, A. (2022). Multiple stressor effects of a heatwave and a herbicide on zooplankton communities: implications of global climate change. Frontiers in Environmental Science, 2212.

Shen, C., Pan, X., Wu, X., Xu, J., Dong, F., & Zheng, Y. (2022). Ecological risk assessment for difenoconazole in aquatic ecosystems using a web-based interspecies correlation estimation (ICE)-species sensitivity distribution (SSD) model. Chemosphere, 289(October 2021), 133236. https://doi.org/10.1016/j.chemosphere.2021.133236.

Silva, V., Yang, X., Fleskens, L., Ritsema, C. J., & Geissen, V. (2022). Environmental and human health at risk – Scenarios to achieve the Farm to Fork 50% pesticide reduction goals. Environment International, l65(January), 107296. https://doi.org/10.1016/j.envint.2022.107296.

Soria, J. M. (2006). Past, present and future of la Albufera of Valencia Natural Park. Limnetica, 25(1-2), 135–142. https://doi.org/10.23818/limn.25.10.

van Wijngaarden, R. P. A., Belgers, D. J. M., Zafar, M. I., Matser, A. M., Boerwinkel, M. C., & Arts, G. H. P. (2014). Chronic aquatic effect assessment for the fungicide azoxystrobin. Environmental Toxicology and Chemistry, 33(12), 2775–2785. https://doi.org/10.1002/etc.2739.

Vera-Herrera, L., Soria, J., Pérez, J., & Romo, S. (2021). Long-term hydrological regime monitoring of a mediterranean agro-ecological wetland using landsat imagery: Correlation with the water renewal rate of a shallow lake. Hydrology, 8(4), 21. https://doi.org/10.3390/hydrology8040172.

Vilas-Boas, J. A., Arenas-Sánchez, A., Vighi, M., Romo, S., Van den Brink, P. J., Pedroso Dias, R. J., & Rico, A. (2021). Multiple stressors in Mediterranean coastal wetland ecosystems: Influence of salinity and an insecticide on zooplankton communities under different temperature conditions. Chemosphere, 269. https://doi.org/10.1016/j.chemosphere.2020.129381.

Willming, M. M., & Maul, J. D. (2016). Direct and indirect toxicity of the fungicide pyraclostrobin to Hyalella azteca and effects on leaf processing under realistic daily temperature regimes. Environmental Pollution, 211, 435–442. https://doi.org/10.1016/j.envpol.2015.11.029.

