## Supplementary Material for "A Bayesian network approach to assess the influence of climate change and pesticide use practices on the ecological risks of pesticides in a protected Mediterranean wetland"

**Text S1. Water balance calculations**

The water balance in each rice plot was estimated from the hydrological management of the crops and the approximate water depth in the field plots (10 cm). The amount of irrigation in each day was calculated as:

IRRIGATION =(10/R)*IRRI + (ETP + SEEP)

where IRRI is a binary value that indicates if the input of water is occurring (IRRI=1) or not (IRRI=0), R is the residence time of water in the rice plot (d), ETP the total evapotranspiration per day (cm/d), and SEEP the seepage rate (cm/d).

The amount of drainage in each day was calculated as:

DRAINAGE =(10/R)*DRAIN + PP

where DRAIN is a binary value that indicates if the drainage of the crop is occurring (DRAIN=1) or not (DRAIN=0), and PP is the total precipitation per day (cm/d).

In this way, the water depth was adjusted daily as:

MODELLED DEPTH = ((IRRIGATION + PP) – (SEEP + ETP + DRAINAGE)) + DEPTH*_t-1_*

where DEPTH*_t-1_* is the modelled depth of the previous day.

**Text S2. Renewal rate calculations**

The water renewal rates of each cluster were calculated considering a measured flow dataset for 2008 from the 62 sampling points (J. Soria, Pers. Comm., 2021), the area of each single cluster and the observed depth in paddy fields. For each cluster:

Flow rate (m^3^/day) = flow measure * 86400

where flow measure is the mean of repeated measures in each point of the dataset between the months of May and October (m^3^/s).

Then, it was calculated the total water volume that a cluster could contain to have a fixed water depth of 10 cm:

Cluster capacity (m^3^) = Area * Paddy depth

where area is the total surface of each cluster (m^2^), and paddy depth is equal to 0.1m.

Finally, the renewal rate of each cluster was calculated as the quotient between the cluster capacity and the flow rate:

Renewal rate (days) = Cluster capacity (m^3^) /Flow rate (m^3^/day)

**Text S3. Evapotranspiration calculations**

For each day, the evapotranspiration (ET*_0_*) was calculated according to the formula described by Hargreaves & Samani (1985):

ET*_0_* = 0.0135 (t*_mean_* + 17.78) R*_s_*

where t*_mean_* is the mean daily temperature, and R*_s_* is the incident solar radiation, that was calculated according to Samani (2000):

R*_s_* = R*_0_* * KT * (t*_max_* - t*_min_*)*^0.5^*

where R*_0_* is the extraterrestrial solar radiation, KT is an empirical coefficient, t*_max_* is the maximum daily temperature, and t*_min_* is the minimum daily temperature.

The value of the empirical coefficient of KT was fixed at 0.19, since we are working with a coastal scenario (Hargreaves & Samani, 1985). The values of R*_0_* were taken from meteorological stations of Spanish Ministry of Agriculture, Livestock, Fisheries and Food (MAPA, 2021). The rest of the values, including the mean daily temperature and total precipitation were taken from the Valencian Association of Meteorology (AVAMET, 2021). Additionally, the conversion of R*_0_* from MJ/m^2^/day to mm/day was calculated as:

R*_0_* (mm/day) = R*_0_* (MJ/m^2^/day)* (238.5 / (597.3 -0.57 t*_mean_*))

**Table S1**. Rice-crop scenario parameters used for the RICEWQ simulations. 1: Dates depending on the assigned cases; 2: Not necessary since each set of runs have the same meteorological data (2008, 2050 or 2100); 3: Default data; 4: Assumption; 5: Field measurement; 6: Depending on hydrological clusters

| RICEWQ parameters | Value |
| --- | --- |
| Begin simulation | Day 1^1^ |
| End simulation | Day 142^1^ |
| Meteo code | (Empty) ^2^ |
| EXAMS Flag (0 = not create EXAMSII transfer file; 1 = create EXAMSII transfer file) | 0^3^ |
| Crop emergence | Day 9^1^ |
| Crop maturation | Day 135 ^1^ |
| Crop harvest | Day 142^1^ |
| Maximum areal coverage of crop (fraction) | 0.50^4^ |
| Deposition of pesticides residues at harvest (-1 = left alone; -2 = foliar residues removed from system) | -1^3^ |
| Surface area of paddy (ha) | Depending on Clusters^6^ |
| Depth of paddy outlet or berm height (cm) | 21.00^5^ |
| Initial depth of water in paddy (cm) | 0.00^5^ |
| Seepage rate (cm/ha/day) | 0.00^4^ |
| Depth of active sediment layer (cm) | 5.00^3^ |
| Field capacity (cm/cm) | 0.35^3^ |
| Wilting point (cm/cm) | 0.24^3^ |
| Initial soil moisture (cm/cm) | 0.35^3^ |
| Bulk density of bed sediment (g/cc) | 1.50^3^ |
| Suspended sediment concentration (mg/L) | 50^3^ |
| EXAMSII environment catalog number | 0^3^ |
| EXAMSII chemical catalog number for chemical 1 | 0^3^ |
| Number corresponding to parent of chemical 2 | 0^3^ |
| Number corresponding to parent of chemical 3 | 0^3^ |
| EXAMSII chemical catalog number for chemical 2 | 0^3^ |
| Signals the type of process transforming from parent to metabolite | 0^3^ |
| Gives the reactive molecular form from parent to metabolite | 0^3^ |
| Product yield from the transformation pathway dimensions of mole of transformation product produced per mole of parent compound reacted | 0^3^ |
| EXAMSII chemical catalog number for chemical 3 | 0^3^ |
| Signals the type of process transforming from parent to metabolite | 0^3^ |
| Gives the reactive molecular form from parent to metabolite | 0^3^ |
| Product yield from the transformation pathway dimensions of mole of transformation product produced per mole of parent compound reacted | 0^3^ |

**Table S2.** Physico-chemical properties for the pesticides included in this study. Reference: 1: Obtained from manufacturers recommendations (L. Blanch, Pers. Comm, 2021); 2: Obtained from Rico *et al.*, (2018); 3: Pesticides Properties Database (Lewis et al. 2016); 4: (European Food Safety Authority (EFSA), 2010); 5: calculated value based on arithmetic mean foliar degradation for all other pesticides; 6: (European Food Safety Authority (EFSA), 2015)

| Parameter | Acetamiprid | Azoxystrobin | Bentazon | Cyhalofop-butyl | Difenoconazole | Imazamox | MCPA | Penoxsulam | Propanil |
| --- | --- | --- | --- | --- | --- | --- | --- | --- | --- |
| Nº applications | 1^1^ | 2^1^ | 1^1^ | 2^1^ | 2^1^ | 1^1^ | 1^1^ | 1^1^ | 1^1^ |
| App rate (kg/ha) | 0.03^1^ | 0.2^1^ | 1^1^ | 0.3^1^ | 0.13^1^ | 0.04^1^ | 0.5^1^ | 0.04^1^ | 0.48^1^ |
| Degradation rate in water (1/d) | 0.04^2^ | 0.114^3^ | 0.0087^3^ | 0.126^3^ | 0.231^3^ | 0.005^3^ | 0.051^3^ | 0.046^6^ | 0.577^3^ |
| Degradation rate in saturated soil (1/d) | 0.231^3^ | 0.003^3^ | 0.09^3^ | 0.976^3^ | 0.231^3^ | 0.003^3^ | 0.040^3^ | 0.03^6^ | 0.577^3^ |
| Degradation rate in unsaturated soil (1/d) | 0.231^3^ | 0.003^3^ | 0.09^3^ | 0.976^3^ | 0.007^3^ | 0.042^3^ | 0.028^3^ | 0.117^6^ | 1.732^3^ |
| Degradation rate on foliage (1/d) | 0.110^3^ | 0.09^3^ | 0.462^3^ | 0.161^3^ | 0.094^3^ | 0.124^5^ | 0.165^3^ | 0.248^3^ | 0.018^3^ |
| Water/sediment partition coefficient (cc/g) | 5.8^3^ | 20.822^4^ | 1.6^3^ | 5.39^3^ | 109.04^3^ | 0.34^3^ | 0.027^3^ | 2.12^3^ | 4.32^3^ |
| Volatilization coefficient (m/day) | 1.1E-05^3^ | 1.12E-06^3^ | 0.014^3^ | 3.89E-07^3^ | 1.4E-04^3^ | 8.6E-10^3^ | 0.011^3^ | 4.1E-12^3^ | 0.087^3^ |
| Solubility in water (ppm) | 2950^3^ | 6.7^3^ | 7112^3^ | 180^3^ | 15^3^ | 626000^3^ | 29390^3^ | 408^3^ | 95^3^ |

**Table S3**. RICEWQ parameter values for pesticide applications. 1:Field observation; 2: Default value; 3: Assumption; 4: European Food Safety Authority (EFSA, 2008); 5: PPDB (Lewis et al. 2016).

| RICEWQ parameters | Value |
| --- | --- |
| Date pesticide is applied | Depending on Cases^1^ |
| Depth of incorporation (cm) | 0.00^2^ |
| Application efficiency (fraction) | 1.00^2^ |
| Drift (%)- used in EXAMSII transfer files | 0.0^2^ |
| Number of chemicals in simulation | 1^3^ |
| Number of transformation paths for simulating metabolites | 0^3^ |
| Flag for temperature dependent biolysis | 1^2^ |
| Pesticide concentration in water (mg/L) | 0.0^2^ |
| Pesticide concentration in benthic sediments (mg/kg) | 0.0^2^ |
| Pesticide concentration in foliage (mg/L) | 0.0^2^ |
| Hydrolysis degradation rate in water (1/day) | 0.0^3^ |
| Photolysis degradation rate in water (1/day) | 0.0^3^ |
| Wash-off rate per cm of precipitation | 0.2^2^ |
| Settling velocity (m/day) | 2.0^2^ |
| Mixing depth to allow direct partitioning to bed (cm) | 0.1^2^ |
| Mixing velocity (diffusion)(m/day) | 0.001^2^ |
| Release rate for slow-release formulation (1/day) | 0.00^2^ |
| Fraction of non-intercepted chemical immediately lost | 0.0^2^ |
| Flag for bi-phase transformation from parent to metabolite | 0^2^ |
| Q10 coefficient for degradation in water | 2.58^4^ |
| Q10 coefficient for degradation in wet sediment | 2.58^4^ |
| Q10 coefficient for degradation in dry sediment | 2.58^4^ |
| Reference temperature for KWM in card 10B | 20^5^ |
| Reference temperature for KSW in card 10B | 20^5^ |
| Reference temperature for KSD in card 10B | 20^5^ |

**Table S4**. Exposure distribution parameters for the pesticide exposure data obtained with the RICEWQ model and for the Species Sensitivity Distributions (SSDs). Ac: acute toxicity; Chr: chronic toxicity; PEC: Peak exposure concentrations; TWAC: Time weighted average concentrations.

| **Compound** | **Scenarios** | **Exposure distribution** | | **Effect distribution** | |
| --- | --- | --- | --- | --- | --- |
|  |  | **Type** | **Parameters** | **Type** | **Parameters** |
| **Acetamiprid** | PEC-2008- | Log-normal | meanlog:-2.3; sdlog: 0.68 | Log-normal | Ac: 5.6 (meanlog), 3.6 (sdlog); Chr: 5.8 (meanlog), 4(sdlog) |
|  | PEC-2008 | Log-normal | meanlog: -1.3 ; sdlog: 0.68 |  |  |
|  | PEC-2008+ | Log-normal | meanlog: -1.01; sdlog: 0.68 |  |  |
|  | PEC-2050- | Log-normal | meanlog: -1.4; sdlog: 0.57 |  |  |
|  | PEC-2050 | Log-normal | meanlog: -0.3; sdlog: 0.57 |  |  |
|  | PEC-2050+ | Log-normal | meanlog: -0.017; sdlog: 0.57 |  |  |
|  | PEC-2100- | Log-normal | meanlog: -1.7; sdlog: 0.34 |  |  |
|  | PEC-2100 | Log-normal | meanlog: -0.6; sdlog: 0.34 |  |  |
|  | PEC-2100+ | Log-normal | meanlog: -0.31; sdlog: 0.34 |  |  |
|  | TWAC-2008- | Log-normal | meanlog: -3.5; sdlog: 0.46 |  |  |
|  | TWAC-2008 | Log-normal | meanlog: -2.4; sdlog: 0.46 |  |  |
|  | TWAC-2008+ | Log-normal | meanlog: -2.11; sdlog: 0.46 |  |  |
|  | TWAC-2050- | Log-normal | meanlog: -2.8; sdlog: 0.43 |  |  |
|  | TWAC-2050 | Log-normal | meanlog: -1.7; sdlog: 0.43 |  |  |
|  | TWAC-2050+ | Log-normal | meanlog: -1.4; sdlog: 0.43 |  |  |
|  | TWAC-2100- | Log-normal | meanlog: -3.2; sdlog: 0.33 |  |  |
|  | TWAC-2100 | Log-normal | meanlog: -2.1; sdlog: 0.3 |  |  |
|  | TWAC-2100+ | Log-normal | meanlog: -1.8; sdlog: 0.33 |  |  |
| **Azoxystrobin** | PEC-2008- | Weibull | shape: 9.9; scale: 42.3 | Log-normal | Ac: 6.3 (meanlog); 1.5 (sdlog); Chr: 3.8 (meanlog); 3 (sdlog) |
|  | PEC-2008 | Weibull | shape: 9.9; scale: 84.7 |  |  |
|  | PEC-2008+ | Weibull | shape: 9.9; scale: 127.1 |  |  |
|  | PEC-2050- | Weibull | shape: 8.6; scale: 38 |  |  |
|  | PEC-2050 | Weibull | shape: 8.6; scale: 76.1 |  |  |
|  | PEC-2050+ | Weibull | shape: 8.6; scale: 114.2 |  |  |
|  | PEC-2100- | Weibull | shape: 9.2; scale: 36.7 |  |  |
|  | PEC-2100 | Weibull | shape: 9.3; scale: 73.4 |  |  |
|  | PEC-2100+ | Weibull | shape: 9.3; scale: 110.1 |  |  |
|  | TWAC-2008- | Weibull | shape: 4.9; scale: 9.1 |  |  |
|  | TWAC-2008 | Weibull | shape: 4.9; scale: 18.3 |  |  |
|  | TWAC-2008+ | Weibull | shape: 4.9; scale: 27.5 |  |  |
|  | TWAC-2050- | Weibull | shape: 5.8; scale: 7.7 |  |  |
|  | TWAC-2050 | Weibull | shape: 5.8; scale: 15.5 |  |  |
|  | TWAC-2050+ | Weibull | shape: 5.8; scale: 23.3 |  |  |
|  | TWAC-2100- | Weibull | shape: 6.3; scale: 7.6 |  |  |
|  | TWAC-2100 | Weibull | shape: 6.3; scale: 15.2 |  |  |
|  | TWAC-2100+ | Weibull | shape: 6.3; scale: 22.8 |  |  |
| **Bentazon** | PEC-2008- | Log-normal | meanlog: 1.7; sdlog: 0.21 | Log-normal | Ac: 9.1 (meanlog); 2.7 (sdlog); Chr: 7 (meanlog); 2.1 (sdlog) |
|  | PEC-2008 | Log-normal | meanlog: 2.4; sdlog: 0.21 |  |  |
|  | PEC-2008+ | Log-normal | meanlog: 2.8; sdlog:0.21 |  |  |
|  | PEC-2050- | Log-normal | meanlog: 1.6; sdlog: 0.33 |  |  |
|  | PEC-2050 | Log-normal | meanlog: 2.3; sdlog: 0.33 |  |  |
|  | PEC-2050+ | Log-normal | meanlog: 2.7; sdlog:0.33 |  |  |
|  | PEC-2100- | Log-normal | meanlog: 1.1; sdlog: 0.44 |  |  |
|  | PEC-2100 | Log-normal | meanlog: 1.7; sdlog: 0.44 |  |  |
|  | PEC-2100+ | Log-normal | meanlog: 2.2; sdlog: 0.44 |  |  |
|  | TWAC-2008- | Log-normal | meanlog: 1.1; sdlog: 0.35 |  |  |
|  | TWAC-2008 | Log-normal | meanlog: 1.8; sdlog: 0.35 |  |  |
|  | TWAC-2008+ | Log-normal | meanlog: 2.2; sdlog: 0.35 |  |  |
|  | TWAC-2050- | Log-normal | meanlog: 0.68; sdlog: 0.34. |  |  |
|  | TWAC-2050 | Log-normal | meanlog: 1.3; sdlog:0.34 |  |  |
|  | TWAC-2050+ | Log-normal | meanlog: 1.7; sdlog: 0.34 |  |  |
|  | TWAC-2100- | Log-normal | meanlog: 0.02; sdlog: 0.41 |  |  |
|  | TWAC-2100 | Log-normal | meanlog: 0.71; sdlog: 0.41 |  |  |
|  | TWAC-2100+ | Log-normal | meanlog: 1.1; sdlog: 0.41 |  |  |
| **Cyhalofop-buthyl** | PEC-2008- | Log-normal | meanlog: 0.53; sdlog: 0.57 | Log-normal | Ac: 7.6 (meanlog); 1.6 (sdlog); Chr: 4.1 (meanlog); 2.7 (sdlog) |
|  | PEC-2008 | Log-normal | meanlog: 1.22; sdlog: 0.57 |  |  |
|  | PEC-2008+ | Log-normal | meanlog: 1.6; sdlog: 0.57 |  |  |
|  | PEC-2050- | Log-normal | meanlog 0.08; sdlog: 0.71 |  |  |
|  | PEC-2050 | Log-normal | meanlog: 0.78; sdlog: 0.71 |  |  |
|  | PEC-2050+ | Log-normal | meanlog: 1.1; sdlog: 0.71 |  |  |
|  | PEC-2100- | Log-normal | meanlog: -1.2; sdlog: 0.33 |  |  |
|  | PEC-2100 | Log-normal | meanlog: -0.57; sdlog: 0.33 |  |  |
|  | PEC-2100+ | Log-normal | meanlog: -0.16; sdlog: 0.33 |  |  |
|  | TWAC-2008- | Log-normal | meanlog: -1.1; sdlog: 0.42 |  |  |
|  | TWAC-2008 | Log-normal | meanlog: -0.48; sdlog: 0.42 |  |  |
|  | TWAC-2008+ | Log-normal | meanlog: -0.08; sdlog: 0.42 |  |  |
|  | TWAC-2050- | Log-normal | meanlog: -1.4; sdlog: 0.51 |  |  |
|  | TWAC-2050 | Log-normal | meanlog: -0.75; sdlog: 0.51 |  |  |
|  | TWAC-2050+ | Log-normal | meanlog: -0.35; sdlog: 0.51 |  |  |
|  | TWAC-2100- | Log-normal | meanlog: -2.6; sdlog: 0.206 |  |  |
|  | TWAC-2100 | Log-normal | meanlog: -1.9; sdlog: 0.205 |  |  |
|  | TWAC-2100+ | Log-normal | meanlog: -1.5; sdlog: 0.205 |  |  |
| **Difenoconazole** | PEC-2008- | Log-normal | meanlog: 2.1; sdlog: 0.15 | Log-normal | Ac: 5.4 (meanlog); 1.9 (sdlog); Chr: 2.5 (meanlog); 2.2 (sdlog) |
|  | PEC-2008 | Log-normal | meanlog: 2.8; sdlog: 0.15 |  |  |
|  | PEC-2008+ | Log-normal | meanlog: 3.3; sdlog: 0.15 |  |  |
|  | PEC-2050- | Log-normal | meanlog: 1.9; sdlog: 0.17 |  |  |
|  | PEC-2050 | Log-normal | meanlog: 2.6; sdlog: 0.17 |  |  |
|  | PEC-2050+ | Log-normal | meanlog: 3.1; sdlog: 0.17 |  |  |
|  | PEC-2100- | Log-normal | meanlog: 1.93; sdlog: 0.17 |  |  |
|  | PEC-2100 | Log-normal | meanlog: 2.62; sdlog: 0.17 |  |  |
|  | PEC-2100+ | Log-normal | meanlog: 3.08; sdlog: 0.17 |  |  |
|  | TWAC-2008- | Log-normal | meanlog: 0.11; sdlog: 1.6 |  |  |
|  | TWAC-2008 | Log-normal | meanlog: 0.8; sdlog: 0.16 |  |  |
|  | TWAC-2008+ | Log-normal | meanlog: 1.26; sdlog: 0.16 |  |  |
|  | TWAC-2050- | Log-normal | meanlog: -0.03; sdlog: 0.07 |  |  |
|  | TWAC-2050 | Log-normal | meanlog: 0.65; sdlog: 0.079 |  |  |
|  | TWAC-2050+ | Log-normal | meanlog: 1.11; sdlog: 0.079 |  |  |
|  | TWAC-2100- | Log-normal | meanlog: -0.02; sdlog: 0.06 |  |  |
|  | TWAC-2100 | Log-normal | meanlog: 0.67; sdlog: 0.06 |  |  |
|  | TWAC-2100+ | Log-normal | meanlog: 1.13; sdlog: 0.06 |  |  |
| **Imazamox** | PEC-2008- | Log-normal | meanlog: 0.62; sdlog: 0.3 | Log-normal | Ac: 6.7 (meanlog); 3 (sdlog); Chr: 3.8 (meanlog); 2.2 (sdlog) |
|  | PEC-2008 | Log-normal | meanlog: 0.91; sdlog: 0.3 |  |  |
|  | PEC-2008+ | Log-normal | meanlog: 1.3; sdlog: 0.302 |  |  |
|  | PEC-2050- | Log-normal | meanlog: 0.73; sdlog: 0.21 |  |  |
|  | PEC-2050 | Log-normal | meanlog: 1.02; sdlog: 0.21 |  |  |
|  | PEC-2050+ | Log-normal | meanlog: 1.42; sdlog: 0.21 |  |  |
|  | PEC-2100- | Log-normal | meanlog: 0.64; sdlog: 0.24 |  |  |
|  | PEC-2100 | Log-normal | meanlog: 0.93; sdlog: 0.24 |  |  |
|  | PEC-2100+ | Log-normal | meanlog: 1.34; sdlog: 0.24 |  |  |
|  | TWAC-2008- | Log-normal | meanlog: 0.44; sdlog: 0.38 |  |  |
|  | TWAC-2008 | Log-normal | meanlog: 0.72; sdlog: 0.38 |  |  |
|  | TWAC-2008+ | Log-normal | meanlog: 1.13; sdlog: 0.38 |  |  |
|  | TWAC-2050- | Log-normal | meanlog: 0.41; sdlog: 0.34 |  |  |
|  | TWAC-2050 | Log-normal | meanlog: 0.7; sdlog: 0.34 |  |  |
|  | TWAC-2050+ | Log-normal | meanlog: 1.11; sdlog: 0.34 |  |  |
|  | TWAC-2100- | Log-normal | meanlog: 0.38; sdlog: 0.35 |  |  |
|  | TWAC-2100 | Log-normal | meanlog: 0.66; sdlog: 0.35 |  |  |
|  | TWAC-2100+ | Log-normal | meanlog: 1.07; sdlog: 0.35 |  |  |
| **MCPA** | PEC-2008- | Log-normal | meanlog: 2.9; sdlog: 0.14 | Log-normal | Ac: 9.9 (meanlog); 2.9 (sdlog); Chr: 6.6 (meanlog); 3 (sdlog) |
|  | PEC-2008 | Log-normal | meanlog: 3.6; sdlog: 0.15 |  |  |
|  | PEC-2008+ | Log-normal | meanlog: 4.02; sdlog: 0.15 |  |  |
|  | PEC-2050- | Log-normal | meanlog: 2.7; sdlog: 0.15 |  |  |
|  | PEC-2050 | Log-normal | meanlog: 3.4; sdlog: 0.15 |  |  |
|  | PEC-2050+ | Log-normal | meanlog: 3.8; sdlog: 0.15 |  |  |
|  | PEC-2100- | Log-normal | meanlog: 2.4; sdlog: 0.18 |  |  |
|  | PEC-2100 | Log-normal | meanlog: 3.1; sdlog: 0.19 |  |  |
|  | PEC-2100+ | Log-normal | meanlog: 3.5; sdlog: 0.19 |  |  |
|  | TWAC-2008- | Weibull | shape: 3.7; scale: 13 |  |  |
|  | TWAC-2008 | Weibull | shape: 3.7; scale: 26.4 |  |  |
|  | TWAC-2008+ | Weibull | shape: 3.7; scale: 39.6 |  |  |
|  | TWAC-2050- | Weibull | shape: 4.4; scale: 8.9 |  |  |
|  | TWAC-2050 | Weibull | shape: 4.4; scale: 17.8 |  |  |
|  | TWAC-2050+ | Weibull | shape: 4.4; scale: 26.7 |  |  |
|  | TWAC-2100- | Weibull | shape: 4.7; scale: 5.5 |  |  |
|  | TWAC-2100 | Weibull | shape: 4.7; scale: 11.1 |  |  |
|  | TWAC-2100+ | Weibull | shape: 4.7; scale: 16.7 |  |  |
| **Penoxsulam** | PEC-2008- | Log-normal | meanlog: -0.86; sdlog: 0.27 | Log-normal | Ac: 7.1 (meanlog); 3.8 (sdlog);Chr: 4.5 (meanlog); 3.4 (sdlog) |
|  | PEC-2008 | Log-normal | meanlog: -0.17; sdlog: 0.28 |  |  |
|  | PEC-2008+ | Log-normal | meanlog: 0.24; sdlog: 0.28 |  |  |
|  | PEC-2050- | Log-normal | meanlog: -0.99; sdlog: 0.27 |  |  |
|  | PEC-2050 | Log-normal | meanlog: -0.3; sdlog:0.27 |  |  |
|  | PEC-2050+ | Log-normal | meanlog: 0.1; sdlog: 0.27 |  |  |
|  | PEC-2100- | Log-normal | meanlog: -1.2; sdlog: 0.14 |  |  |
|  | PEC-2100 | Log-normal | meanlog: -0.6; sdlog: 0.15 |  |  |
|  | PEC-2100+ | Log-normal | meanlog: -0.2; sdlog: 0.15 |  |  |
|  | TWAC-2008- | Weibull | shape: 4.04; scale: 0.301 |  |  |
|  | TWAC-2008 | Weibull | shape: 4.04; scale: 0.6 |  |  |
|  | TWAC-2008+ | Weibull | shape: 4.04; scale: 0.9 |  |  |
|  | TWAC-2050- | Weibull | shape: 3.8; scale: 0.29 |  |  |
|  | TWAC-2050 | Weibull | shape: 3.8; scale: 0.59 |  |  |
|  | TWAC-2050+ | Weibull | shape: 3.8; scale: 0.89 |  |  |
|  | TWAC-2100- | Weibull | shape: 4.2; scale: 0.207 |  |  |
|  | TWAC-2100 | Weibull | shape: 4.2; scale: 0.41 |  |  |
|  | TWAC-2100+ | Weibull | shape: 4.2; scale: 0.62 |  |  |
| **Propanil** | PEC-2008- | Log-normal | meanlog: 0.74; sdlog: 0.27 | Log-normal | Ac: 7.8 (meanlog); 2.1 (sdlog); Chr: 3.4 (meanlog); 1.3 (sdlog) |
|  | PEC-2008 | Log-normal | meanlog: 1.4; sdlog: 0.28 |  |  |
|  | PEC-2008+ | Log-normal | meanlog: 1.8; sdlog: 0.28 |  |  |
|  | PEC-2050- | Log-normal | meanlog: 1.81; sdlog: 0.35 |  |  |
|  | PEC-2050 | Log-normal | meanlog: 2.5; sdlog: 0.35 |  |  |
|  | PEC-2050+ | Log-normal | meanlog: 2.91; sdlog: 0.35 |  |  |
|  | PEC-2100- | Log-normal | meanlog: 0.87; sdlog: 0.35 |  |  |
|  | PEC-2100 | Log-normal | meanlog: 1.56; sdlog: 0.35 |  |  |
|  | PEC-2100+ | Log-normal | meanlog: 1.97; sdlog: 0.35 |  |  |
|  | TWAC-2008- | Normal | mean: 0.31; sd: 0.07 |  |  |
|  | TWAC-2008 | Normal | mean: 0.62; sd: 0.15 |  |  |
|  | TWAC-2008+ | Normal | mean: 0.93; sd: 0.23 |  |  |
|  | TWAC-2050- | Normal | mean: 0.69; sd: 0.15 |  |  |
|  | TWAC-2050 | Normal | mean: 1.39; sd: 0.30 |  |  |
|  | TWAC-2050+ | Normal | mean: 2.08; sd: 0.30 |  |  |
|  | TWAC-2100- | Normal | mean: 0.28; sd: 0.05 |  |  |
|  | TWAC-2100 | Normal | mean: 0.57; sd: 0.12 |  |  |
|  | TWAC-2100+ | Normal | mean: 0.86; sd: 0.18 |  |  |

**Table S5**. Node descriptions of the Bayesian network performed with Netica. DC: discretized continuous, continuous variables were binned into the states. PEC: Peak exposure concentrations; TWAC21: Time weighted average concentrations.

| Node name | Node type | Number of states/Intervals | Node input (equation & assumptions) |
| --- | --- | --- | --- |
| Climate time | Constant | 3 | Scenarios |
| Application scenario | Constant | 3 | Scenarios |
| Scenario combination | Constant | 162 | Combination of scenarios |
| Exposure time | Constant | 2 | PEC or TWAC |
| Endpoint | Constant | 2 | Dependent on Exposure time:  PEC = Acute  TWAC = Chronic |
| Exposure Concentration distribution (ECD) | DC* | 8 | Scenario and Exposure time dependent distribution:  Distribution depending on scenario (see Table S4) |
| Species sensitivity distribution (SSD) | DC | 8 | Endpoint dependent distribution:  Log-normal Distribution (meanlog, sdlog) |
| Risk quotient distribution (RQD) | DC | 4 | RQD = ECD / SSD |

**Figure S1**. Distribution of rice plots and ditch channels with their corresponding flow points in the ANP. The red dots indicate the points where there were water flow data and the green lines indicate the ditch channels in the Natural Park.

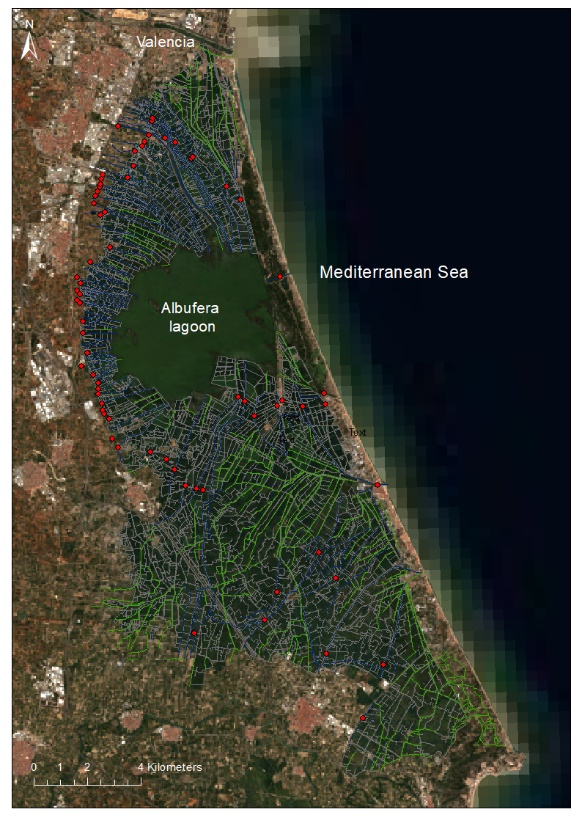

**References**

AVAMET (2021). Valencian Association of Meteorology. Available at: <https://www.avamet.org/>

EFSA, European Food Safety Authority (2008). Opinion on a request from EFSA related to the default Q10 value used to describe the temperature effect on transformation rates of pesticides in soil‐Scientific Opinion of the Panel on Plant Protection Products and their Residues (PPR Panel). *EFSA Journal*, *6*(1), p.622

EFSA, European Food Safety Authority (2010). Conclusion on the peer review of the pesticide risk assessment of the active substance azoxystrobin. *European Food Safety Authority Journal*, *8*(4). https://doi.org/10.2903/j.efsa.2010.1542.

EFSA, European Food Safety Authority (2015). Conclusion on the peer review of the pesticide risk assessment of the active substance penoxsulam. *European Food Safety Authority Journal*, *13(11) 430*(October), 1–90.

Hargreaves, G. H., & Samani, Z. A. (1985). Reference Crop Evapotranspiration From Ambient Air Temperature. *Paper - American Society of Agricultural Engineers*, 96–99.

Lewis, K.A., Tzilivakis, J., Warner, D. and Green, A. (2016) An international database for pesticide risk assessments and management. *Human and Ecological Risk Assessment: An International Journal*, 22(4), 1050-1064. DOI: [10.1080/10807039.2015.1133242](https://doi.org/10.1080/10807039.2015.1133242)

MAPA (2021). Spanish Ministry of Agriculture, Livestock, Fisheries and Food. Meterological stations, available at: https://www.mapa.gob.es/en/

Rico, A., Arenas-Sánchez, A., Pasqualini, J., García-Astillero, A., Cherta, L., Nozal, L. and Vighi, M. (2018). Effects of imidacloprid and a neonicotinoid mixture on aquatic invertebrate communities under Mediterranean conditions. *Aquatic Toxicology*, *204*, pp.130-143.

Samani, Z. (2000). Estimating solar radiation and evapotranspiration using minimum climatological data. *Journal of Irrigation and Drainage Engineering*, 126(4), 265–267.
